# Mitochondrial Complex II In Intestinal Epithelial Cells Regulates T-cell Mediated Immunopathology

**DOI:** 10.1101/2020.03.12.978692

**Authors:** Hideaki Fujiwara, Anna V Mathew, Ilya Kovalenko, Anupama Pal, Ho-Joon Lee, Daniel Peltier, Stephanie Kim, Chen Liu, Katherine Oravecz-Wilson, Lu Li, Yaping Sun, Jaeman Byun, Tom Saunders, Alnawaz Rehemtulla, Costas A Lyssiotis, Subramanian Pennarthur, Pavan Reddy

## Abstract

Intestinal epithelial cell (IEC) damage by T cells contributes to alloimmune, autoimmune and iatrogenic diseases such as graft-versus-host disease (GVHD), inflammatory bowel disease (IBD) and immune checkpoint blockade (ICB) mediated colitis, respectively. Despite significant advances in understanding the aberrant biology of T cells in these diseases, little is known about how the fundamental biological processes of the target IECs influence the disease severity. Here, through analyses of metabolic pathways of IECs, we identified disruption of oxidative phosphorylation without a concomitant change in glycolysis and an increase in succinate levels in several distinct in vivo models of T cell mediated gastrointestinal damage such as GVHD, IBD and ICB mediated colitis. Metabolic flux studies, complemented by imaging and protein analyses identified a critical role for IEC intrinsic succinate dehydrogenase A (SDHA), a component of mitochondrial complex II, in causing these metabolic alterations that contributed to the severity of intestinal damage. The critical mechanistic role of IEC intrinsic SDHA was confirmed by complementary chemical and genetic reduction of SDHA and with IEC specific deletion of SDHA. Further in vitro and in vivo mechanistic studies demonstrated that loss of SDHA in IECs was mediated by the perforin-granzyme from the T cells. The loss of SDHA was also validated in human clinical samples. These data identify a critical role for the alteration of the IEC specific mitochondrial complex II component SDHA in the regulation of the severity of T cell mediated intestinal diseases.

## INTRODUCTION

Intestinal epithelial cells (IECs) function as critical barriers that protect from pathogen invasion. This function is disrupted by environmental factors and/or host genetic factors that cause activation of immune responses that leads to IEC damage and perpetuation of the aberrant inflammation ^1 2 3 4^. The non-infectious inflammatory diseases that occur in the intestinal tract such as autoimmune inflammatory bowel diseases (IBD), immune checkpoint blocker (ICB) mediated colitis, and alloimmune T cell mediated gastrointestinal graft-versus-host disease (GI-GVHD) often have similar symptoms and cause significant morbidity and mortality ^1 5 6^. Although the pathophysiology of each of these diseases is a complex interplay of the environmental and genetic factors, T cell mediated damage of IECs plays a vital role in the cause and severity of all of these conditions. Immunosuppression with corticosteroids, and anti-cytokine therapies have shown good results with adverse effects, but are still incomplete ^5 7 8^. The biology and treatments of these diseases are often understood and targeted from the immune cell and inflammation perspective, but the pathogenesis and severity from host IEC target cell perspective remain undefined.

Emerging data in recent years have brought into focus the central role of immune cell metabolism in the regulation of intestinal inflammatory diseases. APCs, including macrophages and dendritic cells, and T cells show changes in metabolic programming, including morphological alterations in mitochondria, transitioning from glycolysis dependence (pro-inflammatory macrophages and effector T cells) to oxidative phosphorylation (OXPHOS) dependence (anti-inflammatory macrophages and memory T cells) ^9 10 11^ and regulate intestinal T cell mediated diseases such as GI GVHD and IBD. However, whether the metabolism of the targets of these pathogenic T cells in these diseases, the IECs, are perturbed or reprogrammed, and if so, whether this has an impact on the disease severity remains unknown. The mammalian GI tract is a relatively hypoxic region and thus, IECs are uniquely adapted to this hypoxic environment ^12 13^. In this study, we aimed to determine the metabolic changes in IECs when they are targeted by pathogenic T effector cells. We found that in the context of pathogenic T cell mediated damage, the IECs demonstrated a reduction in OXPHOS and accumulated succinate as a result of decrease in SDHA, a component of mitochondrial complex II. The reduction of SDHA in IECs aggravated disease severity in multiple T cell mediated models such as the alloimmune mediated GI GVHD, autoimmune IBD and in ICB mediated colitis. Further mechanistic studies demonstrated that the disruption of mitochondrial complex II by the reduction of SDHA in the IECs was caused by direct engagement and release of perforin dependent granzyme B by the cytotoxic T cells (CTLs) into IECs. Finally the results were validated in human samples of GI GVHD. These results demonstrate that targeting IEC intrinsic metabolism regulates intestinal disease severity mediated by pathogenic T cells.

## RESULTS

### Analysis of mitochondrial respiration in IECs

We explored the impact of immune mediated attack on the bioenergetics of intestinal target cells, IECs. B6 WT animals were lethally irradiated and transplanted with bone marrow (BM) and splenic T cells from either syngeneic B6 or allogeneic BALB/c donors. CD326^+^ IECs from un-transplanted naïve B6 animals, the syngeneic, and allogeneic animals were harvested 21 d after hematopoietic stem cell transplantation (post HCT) and their bio-energetic profiles were analyzed with Seahorse XF ^14^. Compared with IECs from naïve and syngeneic animals, IECs from allogeneic animals demonstrated significantly lower oxygen consumption rates (OCR), but similar extracellular acidification rates (ECAR, an indicator of glycolysis), which dramatically reduced the OCR/ECAR ratio (Fig. 1a-c). IECs from allogeneic animals also did not respond to treatment with carbonyl cyanide-p-trifluoromethoxyphenylhydrazone (FCCP), a mitochondrial uncoupler, when compared to IECs from syngeneic animals (Fig. 1b) demonstrating that the mitochondrial electron transport chain (ETC) functions in IECs from allogeneic mice was reduced. ETC reactions begin at both complex I and complex II. Therefore, to analyze complex I function in IECs, we isolated mitochondria from recipients 21 d post HCT and analyzed their ability to oxidize NADH to NAD^+^. There was no difference in the mitochondrial complex I functions in the IECs harvested from syngeneic and allogeneic recipients (Fig. 1d). Although mitochondrial complex I function was not altered, the net NADH/NAD^+^ ratio was lower in IECs from allogeneic animals than in IECs from syngeneic animals 7 d and 21 d post HCT (Fig. 1e) suggesting that complex II function may be the cause for altered ETC in IECs after allo-HCT. Therefore, we next assessed for mitochondrial complex II and the other super complex levels, with Blue Native Polyacrylamide Gel Electrophoresis (BN-PAGE). Analysis of IECs from naïve, syngeneic, and allogeneic recipients post HCT with BN-PAGE showed alteration of mitochondrial complex II but did not show changes in other mitochondrial supercomplex levels (Fig. 1f and g). These results demonstrated that allo-HCT induced changes in mitochondrial ETC and caused disruption of complex II in IECs.

**Figure 1:**
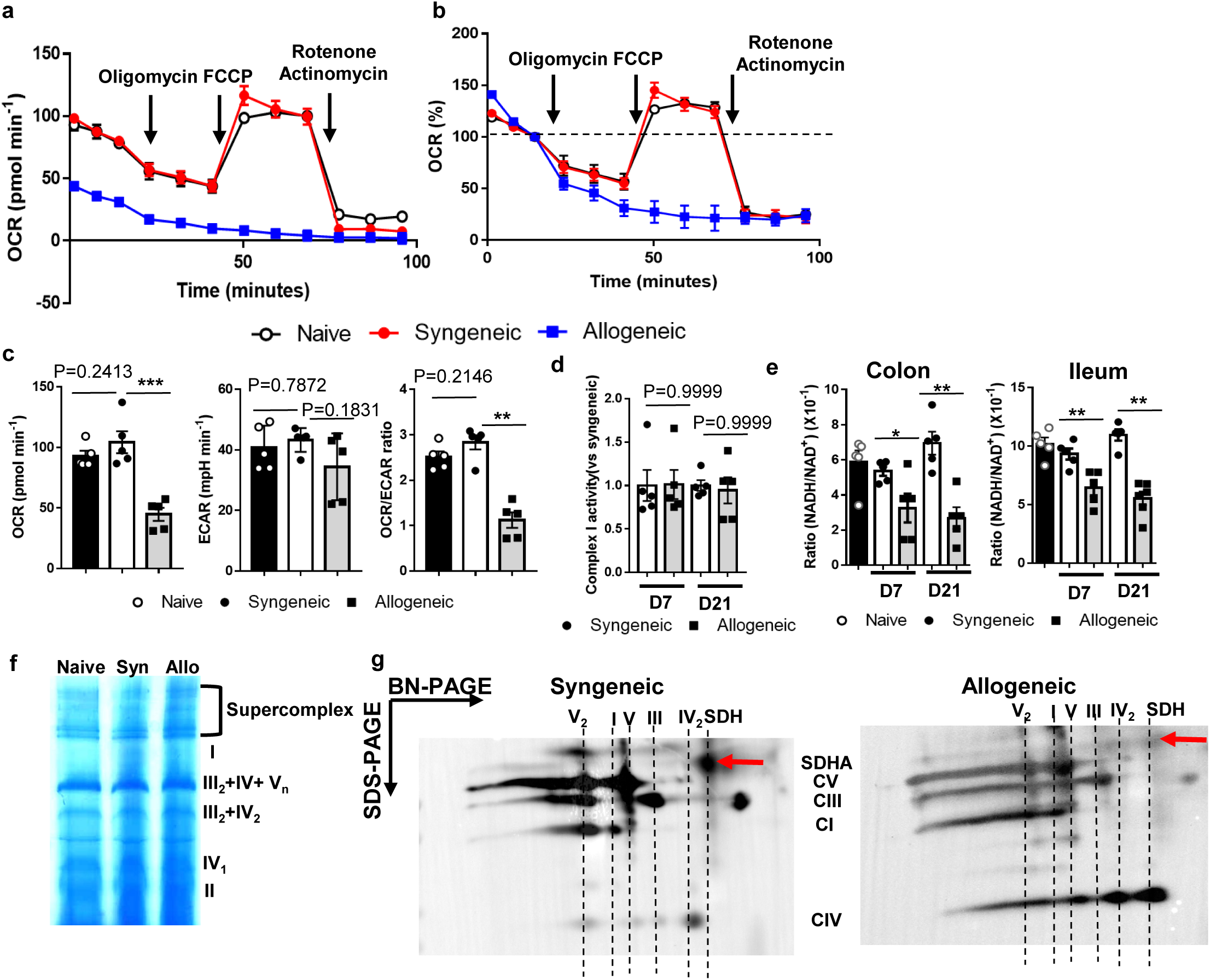
Mitochondrial respiration in IECs change after allo-HCT. B6 WT animals received 10 Gy total body irradiation and received 3×10^6^ T cells along with 5×10^6^ TCD-BM cells from either syngeneic B6 WT or allogeneic BALB/c donors. (**a**) Representative images of SDH enzyme activity staining in GVHD target tissues (colon, ileum, liver and skin) and non-target tissues (heart, pancreas and kidney) from naive animals or recipients 21days post HCT (Scale bar: yellow 500µm, black 200 µm). (**b**) Integrated intensity of SDH enzyme activity staining from colon and ileum 7days post HCT (n=5). Representative plots and a graph summarizing the results of at least two independent experiments are shown. All statistical analysis by Mann-Whitney test (**b**) (mean ± s.e.m.): \**P* < 0.05, \*\**P* < 0.01.

### TCA cycle metabolite profiling in IECs

**Figure 2:**
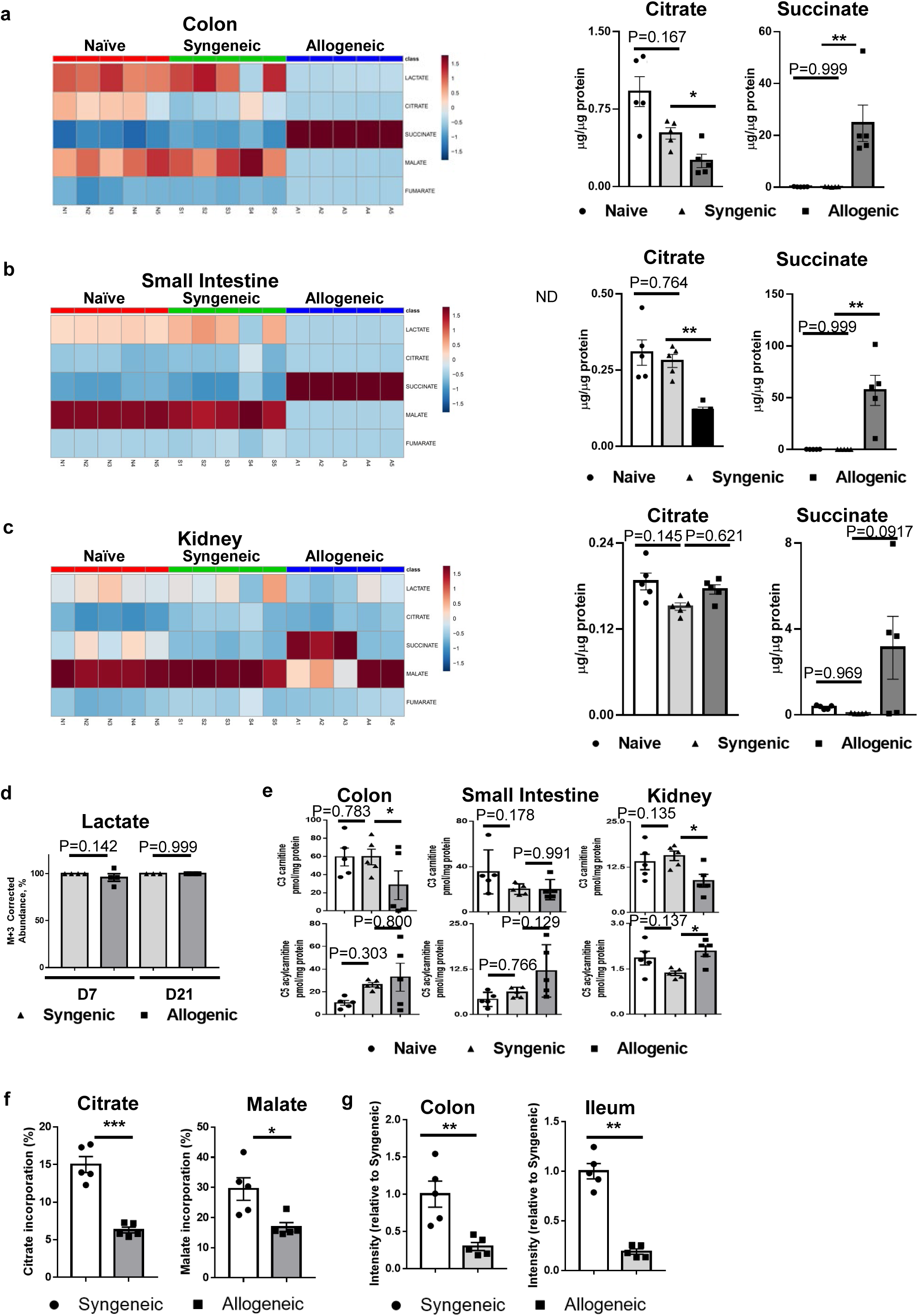
Increased levels of succinate in IECs post-allo-HCT. B6 WT animals received 10 Gy total body irradiation followed by 3×10^6^ T cells along with 5×10^6^ TCD-BM cells from either syngeneic B6 WT or allogeneic BALB/c donors. (**a**) Immunoblot protein density quantification of SDHA and SDHB in mitochondria from IECs 7days post HCT (n=4). (**b**) Representative images of immunofluorescence staining of kidney and liver from recipients 21days post HCT (Complex IV=green, SDHA=red, DAPI=blue, scale bar= 50 µm, n=5). (**c**) Numbers of gold particles per mitochondrial from colon and ileum of naïve animals or recipients 7days post HCT in transmission electron microscopy images with immune-gold staining of SDHA (total 50 mitochondria from 3 samples). (**d**) Representative images of transmission electron microscopy in mitochondria of colon from naive or recipients 21days post HCT (left; scale bar 200nm). Arrow indicates normal cristae and arrowhead indicates abnormal cristae (n=3). (**e**) Mitochondria DNA relative copy numbers of colon and ileum from syngeneic and allogeneic recipients 7 and 21days post HCT (n=5). Representative plots and a graph summarizing the results of at least two independent experiments are shown. All statistical analysis by Mann-Whitney test (**a, e**) or one-way ANOVA analysis with Tukey post hoc test (**c**) (mean ± s.e.m.): \*\*\*\**P* < 0.0001.

### Mechanisms for increase of succinate levels in the IECs

We next explored the mechanisms for succinate accumulation in the IECs from allogeneic recipients. IECs utilize fatty acids supplied from beta-oxidation as their key sources of energy^15, 16^. Therefore, high levels of succinate could be the result of enhanced beta oxidation or anaplerosis. We investigated carnitine levels in each organ as a surrogate for beta-oxidation, and no differences were observed among the groups in the IECs and also in the various organs (Fig. 2e). We next performed metabolic flux analysis (MFA) with LC/MS to assess incorporation of ^13^C-glucose into IEC glucose pools after transplantation. We treated syngeneic and allogeneic transplant recipients with an oral bolus of ^13^C-glucose or ^12^C-glucose (the control) 7 d after the bone marrow transplant (BMT). The IECs from the syngeneic and allogeneic animals were harvested 4 h later and analyzed for incorporation of ^13^C. There was significantly less incorporation of ^13^C into malate in the IECs harvested from allogeneic animals suggesting a reduction in TCA cycle activity (Fig. 2f). To investigate whether anaplerosis from glutamine contributed to succinate accumulation in the TCA cycle we once again performed MFA with LC/MS to measure ^13^C-glutamine in the IECs isolated from various above groups. To this end, IECs were harvested from syngeneic and allogeneic transplant recipients 7 and 21 days after HCT, treated with ^13^C-glutamine or ^12^C-glutamine for 4 h, and then analyzed for incorporation of ^13^C. We detected similar levels of ^13^C incorporation into both alpha-ketoglutarate and succinate in syngeneic and allogeneic IECs (Supplementary Fig. 1) suggesting that enhanced anaplerosis from glutamine did not contribute to greater levels of succinate in the allo-IECs.

### Validation of reduction in SDHA component of mitochondrial complex II

Complex II of mitochondria contains other SDH subunits in addition to SDHA. We therefore next examined SDHA protein levels after HCT to determine whether the SDH dysfunction was due to reduced SDHA protein levels. Mitochondrial proteins were isolated from colonic IECs post HCT and levels of ETC-related proteins (complex I to V) were determined as controls. Reduced SDHA protein levels were observed in IECs from allogeneic animals at 7 d and 21 d post HCT, but no differences were observed in the levels of the other mitochondrial complexes (Figure 3a and Supplementary Fig. 3a).

**Figure 3:**
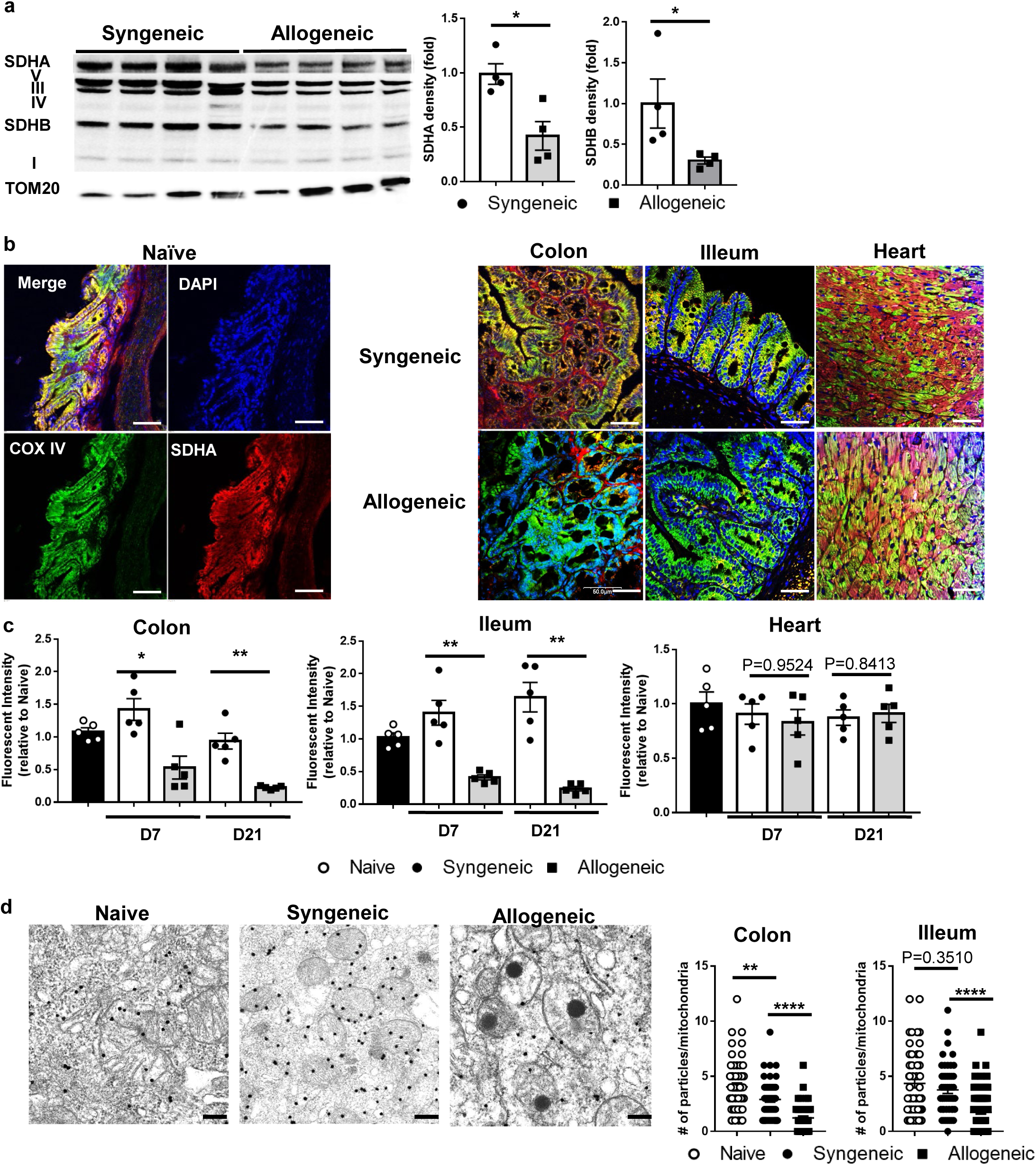
Reduction of SDHA protein levels in IECs post allo-HCT. B6 WT animals received 10 Gy total body irradiation and 3×10^6^ T cells along with 5×10^6^ TCD-BM cells from either syngeneic B6 WT or allogeneic BALB/c donors. (**a**) Immunoblot (left) of SDHA, complex V, III, IV, SDHB, I and TOM-20 in mitochondria from IECs 21days post HCT (n=4), and protein density of SDHA and SDHB (right) (n=4). (f) Representative BN-PAGE image of mitochondria from IECs 21days post HCT showing abundant mitochondrial protein complexes (n=4). (**b**) Representative images of immunofluorescence staining of colon, ileum and heart from naϊve or recipients 21days post HCT (Complex IV=green, SDHA=red, DAPI=blue, scale bar=50 μm, n=5). (**c**) Fluorescent intensity of SDHA in colon, ileum and heart from naϊve, syngeneic or allogeneic recipients 21days post HCT (n=5). (**d**) Representative images of transmission electron microscopy with immune-gold staining for SDHA in mitochondria of colon from naive mice or recipients 21days post HCT (left; scale bar 200nm) and numbers of gold particles per mitochondrial in colon and ileum 21days post HCT (**right**) (50 mitochondria from 3 samples). Representative plots and a graph summarizing the results of at least two independent experiments are shown. Mann-Whitney test (a, c) or one-way ANOVA analysis with Tukey post hoc test (**d**) (mean ± s.e.m.) were used to determine significance: *P < 0.05, **P < 0.01.

### Reduction of SDHA in IECs is specific for T cell mediated intestinal immunopathologies

Next, we first explored whether the reduction in SDHA is germane to any type of intestinal damage, such as conditioning with irradiation or is limited to T cell mediated GVHD in a strain independent manner. To first rule of strain and model dependence in the induction of intestinal GVHD, we next utilized the lethally irradiated major histocompatibility complex (MHC)-matched, multiple host minor histocompatibility antigens (mHAgs)-mismatched C3H.SW→C57BL/6 allogeneic HSCT model of GVHD as in Methods. The IECs harvested from the intestines of the allogeneic recipients showed significantly lower SDHA levels (Supplementary Fig. 4a and b). To determine whether irradiation dependent conditioning was critical, we next utilized well-established chemotherapy-based conditioning regimen consisting of treatment with busulfan and cyclophosphamide. The IECs harvested from allogeneic animals, once again demonstrated significantly lower levels of SDHA (Supplementary Fig. 4c and d). We next analyzed whether allorective T cells alone were sufficient to cause reduction in SDHA independent of conditioning. To this end we utilized the fourth model system, the MHC-haploidentical, non-irradiated parent into F1, the B6→B6D2F1, where alloreactive donor T cells cause GVHD despite absence of any conditioning. Allogeneic IECs once again demonstrated significantly reduced levels of SDHA (Fig. 4a and b). Furthermore, the reduction in SDHA was associated with the observed metabolic abnormality, the accumulation of succinate in the IECs of allogeneic mice (Fig. 4c).

**Figure 4:**
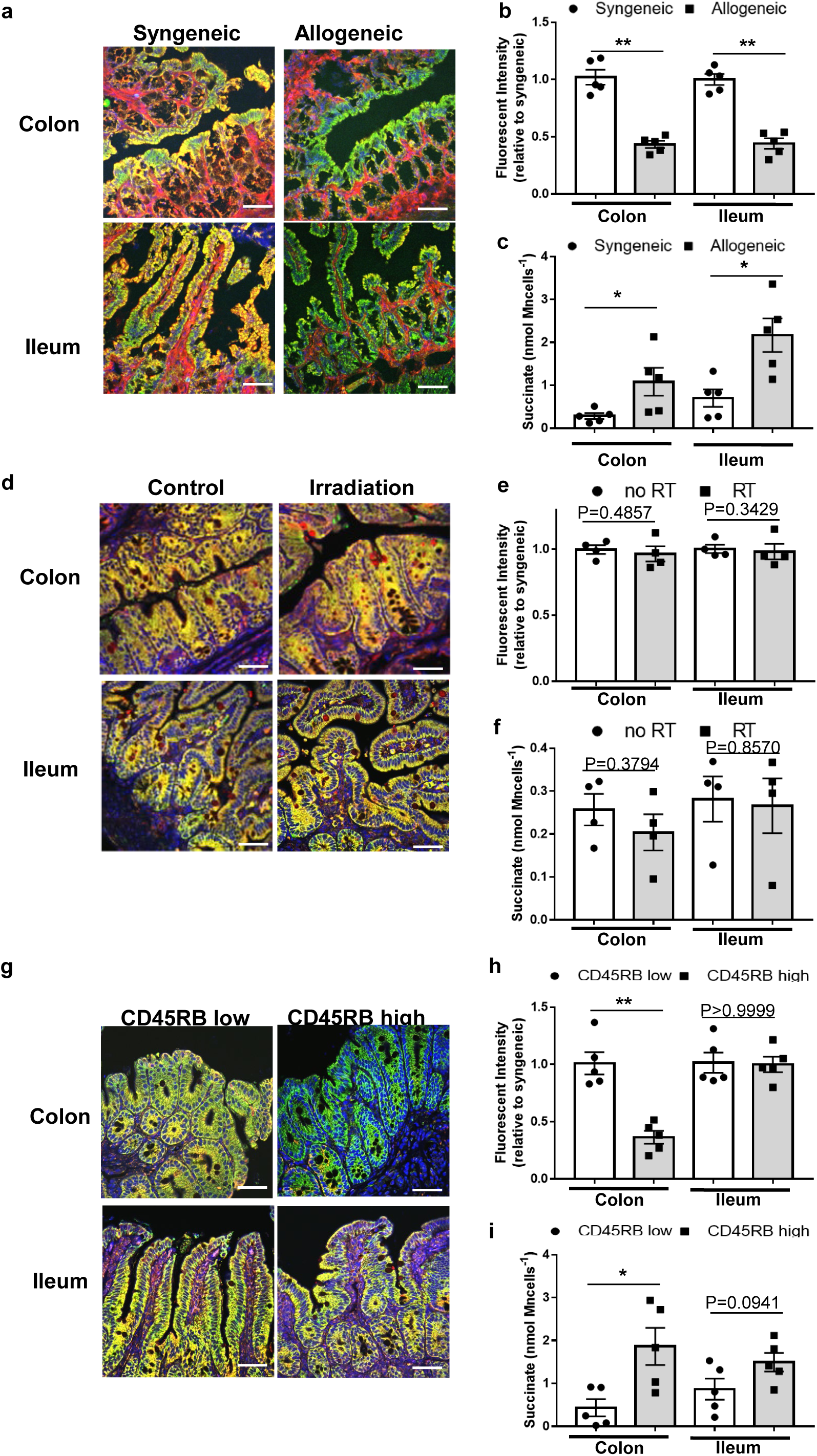
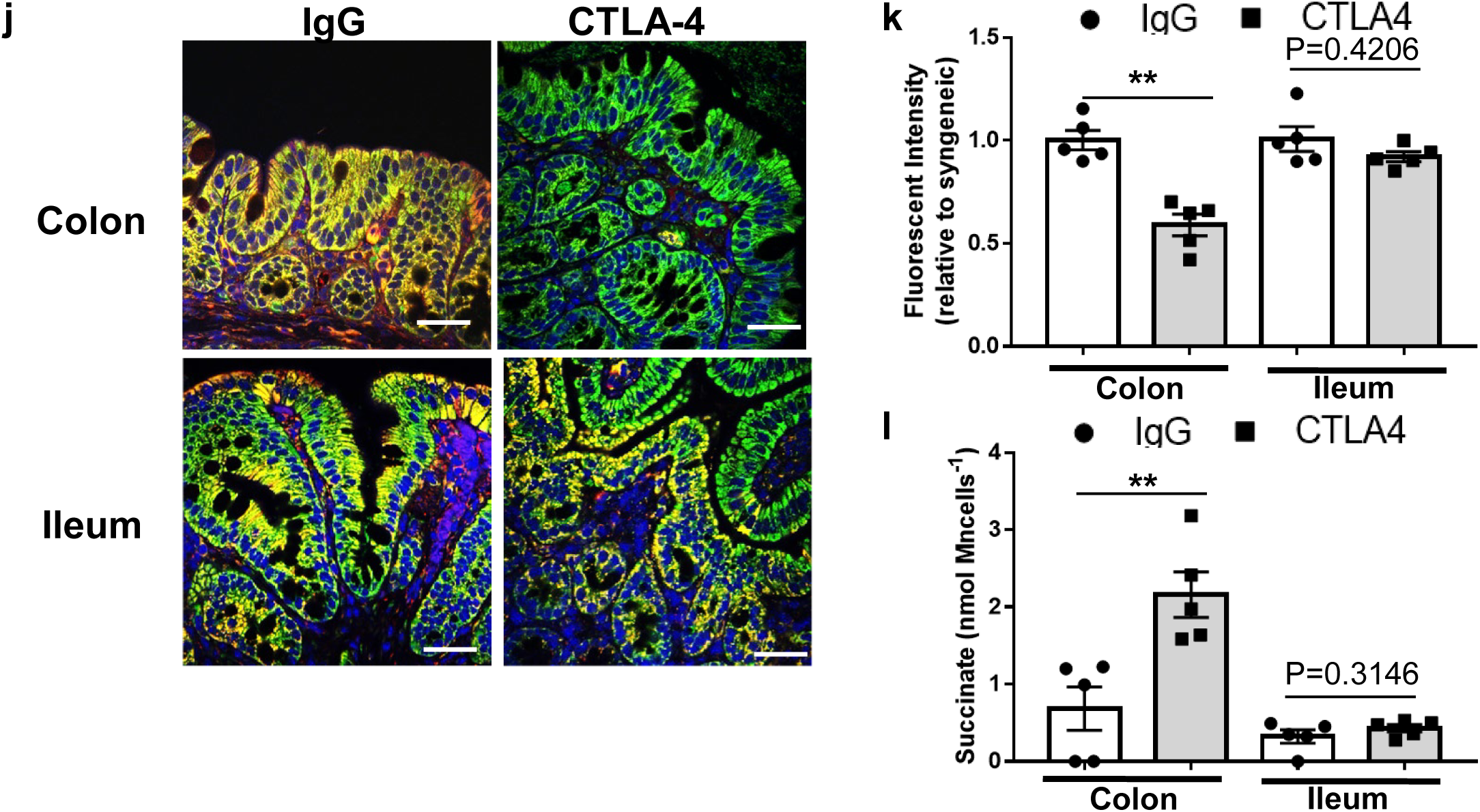
T-cell mediated intestinal immunopathology and SDHA reduction in IECs. (a-c) Unirradiated B6D2F1 mice received 10 × 107 splenocytes from syngeneic B6D2F1 or allogeneic B6 donors (n=5). (**a**) Representative images of immunofluorescence staining and (**b**) fluorescent intensity of SDHA in colon and ileum from B6D2F1 recipients 21days post HCT (scale bar= 50 μm). (**c**) Succinate levels in isolated IECs from colon and ileum 21days post HCT. (d-f) B6 WT mice received 10 Gy total body irradiation without T-cell or BM cells (n=4). (**d**) Representative images of immunofluorescence staining and (**e**) fluorescent intensity of SDHA in colon and ileum from irradiated or non-irradiated (**control**) mice 9days post irradiation (scale bar= 50 μm). (**f**) Succinate levels in isolated IECs from colon and ileum 9days post irradiation. (g-i) CD4+CD25-CD44-CD45RBhi (naϊve) T cells or CD4+CD25-CD44-CD45RBlow (non-naϊve) T cells from B6 WT mice were transferred to RAG-1-/- mice (n=5). (**g**) Representative images of immunofluorescence staining and (**h**) fluorescent intensity of SDHA in colon and ileum 8 weeks after induction of colitis (scale bar= 50 μm). (**i**) Succinate levels in isolated IECs from colon and ileum at 8 weeks after T cell transfer. (j-l) B6 WT mice receiving isotype control IgG or anti-CTLA-4 antibody were treated with 3% DSS in drinking water for 7 days (n=5). (**j**) Representative images of immunofluorescence staining and (**k**) fluorescent intensity of SDHA in colon from mice receiving IgG or CTLA-4 antibody 12days after DSS treatment (scale bar= 50 μm). (**l**) Succinate levels in isolated IECs from colon and ileum 12days after 3% DSS administration. Representative plots and a graph summarizing the results of at least two independent experiments are shown. All statistical analysis by Mann-Whitney test (b, e, h, k) or unpaired t-test (c, f, i, l) (mean ± s.e.m.) were used to determine significance: *P < 0.05, **P < 0.01.

It is possible that any non-T cell mediated damage of the host IECs could also lead to reduction in their SDHA. Therefore, we irradiated mice with escalating doses to cause intestinal damage but without HCT to induce radiation induced colitis. The IECs harvested 7 days after irradiation did not demonstrate a reduction in SDHA levels nor an increase in succinate levels when compared to non-irradiated controls (Fig. 4d-f) suggesting that unlike alloreactive T cell mediated GVHD, radiation induced colitis did not alter SDHA in the IECs.

To further analyze the specificity to T cell mediated damage, we next induced colitis by chemical damage. To this end we next utilized the well-established dextran sulfate sodium (DSS)-chemical induced inflammatory colitis model system. In contrast to autoimmune T cell mediated colitis, twelve days after DSS treatment, the colonic IECs from DSS colitis-induced animals demonstrated similar SDHA and succinate levels as the control animals (Supplementary Fig. 4e-g). These data demonstrate that the specific IEC metabolic abnormality, the reduction in SDHA with accumulation of succinate, was observed exclusively in T cell mediated autoimmune colitis or alloimmune GVHD but not in radiation or chemical colitis.

We further explored whether these observations were broadly applicable to other clinically relevant T cell mediated colitis diseases. T cell dependent immune checkpoint blockers (ICB) have emerged as major form of therapy against many cancers. Amongst the adverse effects of this therapy, the unleashing of T cell mediated colitis is a major toxicity from these therapies, especially with the use of anti-CTLA-4 therapy^18^. Therefore, we next explored whether IECs from ICB mediated T cell colitis demonstrate the aforementioned metabolic abnormality, i.e. reduction in SDHA with an increase in levels of succinate. To this end we utilized the recently developed experimental model for ICB mediated colitis^19, 20^ and administered anti-CTLA-4 antibody before treatment with low dose DSS to induce ICB mediated colitis ^19, 20^. The anti-CTLA-4 treatment induced severe colitis with significant body weight loss, and the colonic IECs demonstrated reduced SDHA with increase in succinate than the control IgG treated animals (Fig. 4j-l and Supplementary Fig. 4h). Collectively, these findings from eight distinct in vivo models demonstrate that T cell mediated intestinal immunopathology cause a reduction in mitochondrial complex II component SDHA with a resultant increase in succinate in their target cells, the IECs.

### Functional relevance of reduction in SDHA in IECs

**Figure 5:**
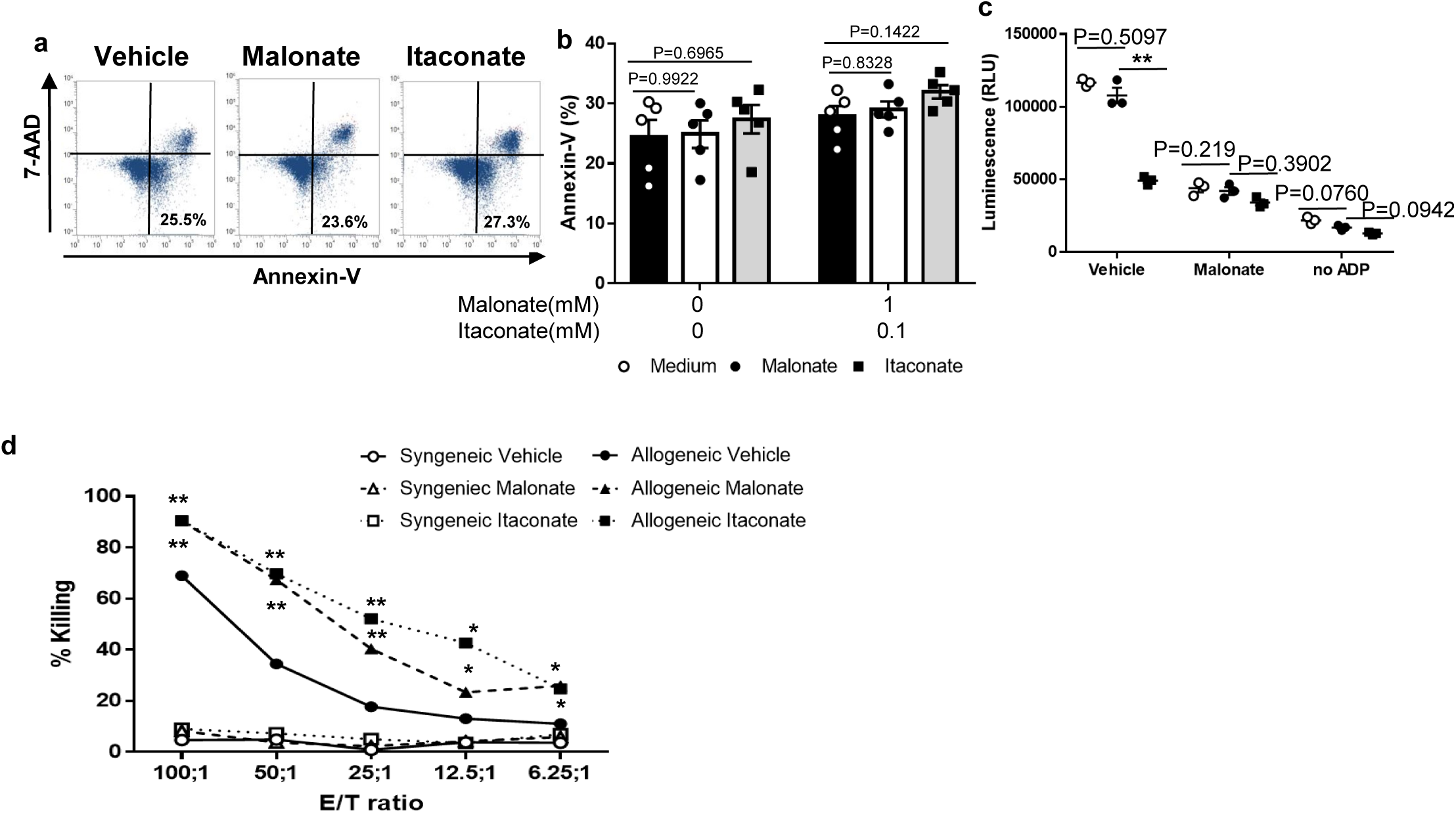
Functional relevance of SDHA reduction in IECs. (a-b) Primary colonic epithelial cells (PCECs) were treated with DMSO, malonate or itaconate for 6 hours (n=5). (**a**) Representative FACS plots and (**b**) Annexin-V positive cells are shown. (**c**) B6 WT animals received 10 Gy total body irradiation and received 3×10^6^ T cells along with 5×10^6^ TCD-BM cells from either syngeneic B6 WT or allogeneic BALB/c donors. ATP production in isolated mitochondria of colon from naϊve, syngeneic and allogeneic recipients 21 days post HCT (n=3). Mitochondria were treated with vehicle, malonate or vehicle without ADP. (**d**) Isolated splenic T cells from either syngeneic B6 WT or allogeneic BALB/c animals were cultured with irradiated splenocytes derived from B6 WT animals for 96 hours, and CD8+ T cells were purified as effector cells. PCECs from B6 WT animals were treated with vehicle, malonate or itaconate and used as targets for effector T-cells. PCECs was analyzed (n=3 biological replicates and n=2 technical replicates). Representative plots and a graph summarizing the results of at least two independent experiments are shown. All statistical analysis by one-way ANOVA analysis with Tukey post hoc test (mean ± s.e.m.) were used to determine significance: *P < 0.05, **P < 0.01, ***P < 0.001.

To functionally demonstrate that inhibition of SDHA leading to ROS generation in the IECs enhanced their sensitivity to T cell mediated apoptosis, B6 PCECs were pretreated with malonate or itaconate at non-apoptosis inducing concentrations and then incubated as targets in a chromium release cytotoxic killing assay with primed T cells. PCECs pre-treated with malonate or itaconate showed significantly higher levels of cell death compared with diluent treated PCECs when co-cultured with allo-primed T cells. In contrast the PCECs that were treated with malonate or itaconate or diluent control and cultured with syngeneic T cells (Fig. 5d) showed no increase in apoptosis. These data demonstrate that reduction or inhibition of SDHA generates mitochondrial ROS in IECs and enhanced their susceptibility T cell mediated killing.

### SDHA in IECs regulates the severity of T cell mediated intestinal immunopathology

Because in vitro data suggested that reduction of SDHA in enterocytes reduced their ability to withstand T cell mediated cytotoxicity, we next examined the role of SDHA in the enterocytes in vivo, in regulating the severity of alloreactive T cell mediated intestinal GVHD. To this end, we took multiple complementary, but distinct approaches. First, we utilized three distinct chemical inhibitors of SDHA, namely malonate, itaconate and atpenin A5. They were administered orally by gavage to naive B6 WT and the IECs from colon and ileum were harvested 12 h after gavage and were stained for SDH enzymatic reactions. The doses at which they inhibited SDHA function 12 h after *in vivo* administration (Supplementary Fig. 6a) were then utilized for analyzing the effect of SDHA inhibition on GI GVHD. Briefly, the recipient animals were gavaged either diluent control or itaconate or malonate or atpenin A5 every other day starting on day 0 post HCT. All of the syngeneic mice survived regardless of treatment, but the allogeneic mice receiving itaconate or malonate or atpenin A5 died significantly faster, and had more severe GVHD symptoms than vehicle treated allogeneic mice (Fig. 6a-b and Supplementary Fig. 6b). Histopathological analysis confirmed increased GVHD severity in the colon and ileum, but not in the liver, skin, and lungs, of the allogeneic recipients (Fig. 6c and Supplementary Fig. 6c).

**Figure 6:**
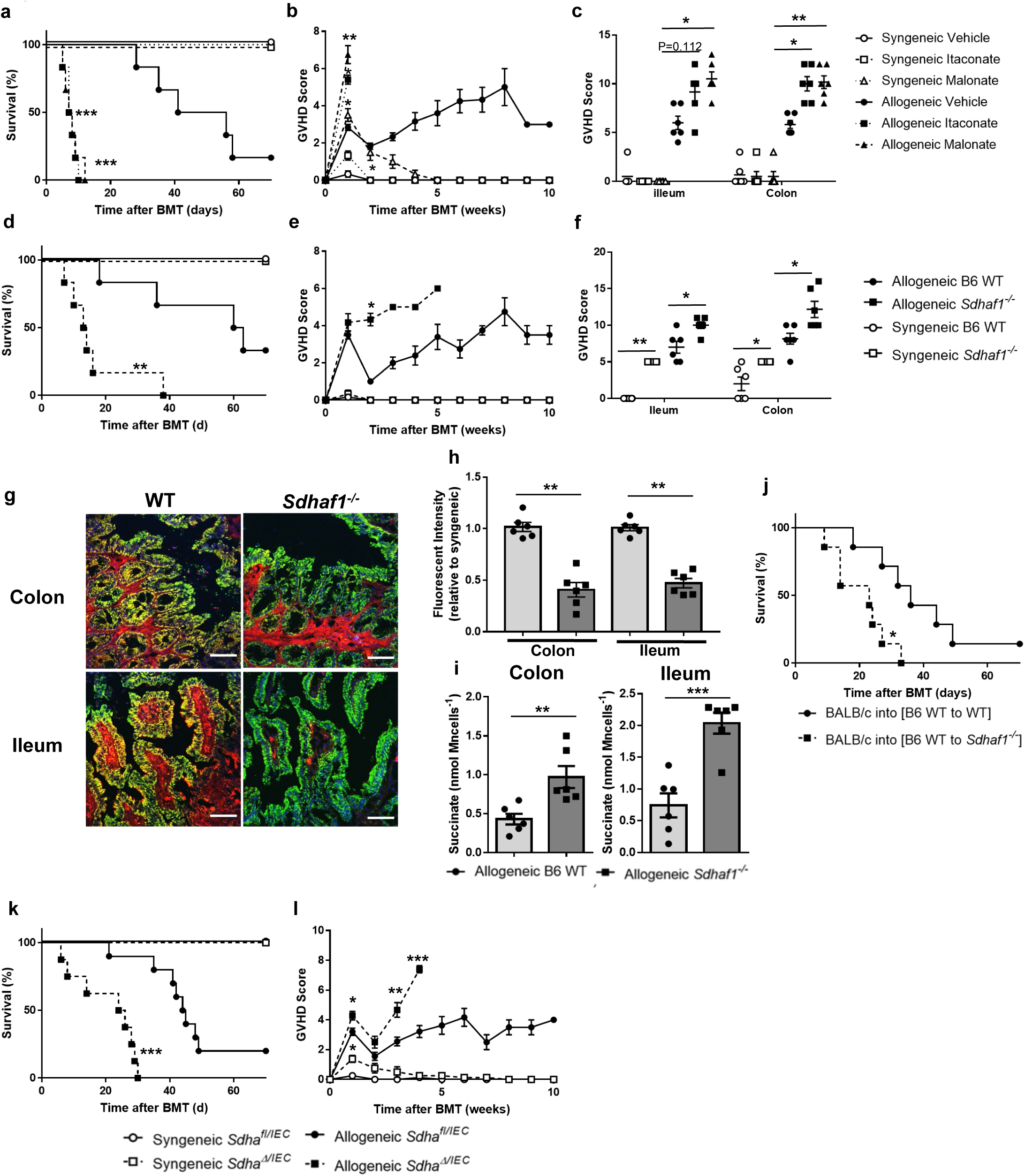
SDHA in IECs regulates the severity of GVHD. B6 WT, Sdhaf1-/-, Sdha? or Sdha animals received 10 Gy total body irradiation followed by 3×10^6^ T cells and 5×10^6^ TCD-BM cells from either syngeneic B6 WT or allogeneic BALB/c donors. (a-c) B6 WT recipients were treated with vehicle, malonate (5g kg-1) or itaconate (2.5g kg-1) every other day post HCT. Survival (**a**), clinical GVHD severity (**b**) and pathological GVHD scores in ileum and colon 7days post HCT (**c**) (n=6). (d-f) B6 WT and Sdhaf1-/- mice received HCT (n=6). Survival (**d**), clinical GVHD severity (**e**) and pathological GVHD scores in ileum and colon 7days post HCT (**f**). (**g**) Representative images of immunofluorescence staining and (**h**) fluorescent intensity of colon and ileum from B6WT and Sdhaf1-/- mice 7days post allo-HCT (Complex IV=green, SDHA=red, DAPI=blue, scale bar= 50 μm, n=6). (**i**) Succinate levels in isolated IECs from colon and ileum of B6WT and Sdhaf1-/-mice at day7 post allo-HCT (n=6). (**j**) Survival of chimeric [B6 Ly5.2? WT] and [B6 Ly5.2 Sdhaf1] animals receiving 9 Gy TBI followed by 3×10^6^ T cells and 5×10^6^ TCD-BM cells from syngeneic B6WT or allogeneic BALB/c donors. (**k**) Survival and clinical GVHD severity (**l**) of Sdha? and Sdha mice post HCT (syngeneic Sdha? and Sdha: n=6, allogeneic Sdha?: n=8, allogeneic Sdha: n=10). Representative plots and a graph summarizing the results of at least two independent experiments are shown. All statistical analysis by log-rank test (a, d, j, k), Mann-Whitney test (b, e, f, h, l), Kruskal-Wallis analysis with Dunn's post hoc test (**c**) or unpaired t-test (**i**) (mean ± s.e.m.): *P < 0.05, **P < 0.01, ***P < 0.001.

It is possible that *Sdhaf1^-/-^* B6 animals might have IECs are more susceptible to any intestinal damage and not uniquely to T cell mediated pathology. To rule out this possibility we induced chemical colitis with DSS in *Sdhaf1^-/-^* B6 animals and compared severity with B6 WT littermates. The *Sdhaf1^-/-^* and the WT animals demonstrated similar changes in body weight and colitis severity demonstrating that the reduction in SDHA predisposed the enterocytes only to T cell mediated cytotoxicity but did not make them more susceptible to non-T cell (i.e. DSS) mediated damage (Supplementary Fig. 6e).

The above approaches, given the systemic effects, do not distinguish potential confounding effects of SDHA activity in host hematopoietic, immune and other cell types from the direct effects on intestinal epithelial target cells. Therefore, first to determine whether SDHA expression exclusively in the host target non-hematopoietic cells is critical for regulation of GI GVHD, we generated BM chimeric mice wherein low SDH activity was restricted to the non-hematopoietic recipient cells. The [WT B6Ly5.2→ WT B6] and [WT B6Ly5.2→ *Sdhaf1^-/-^* B6] chimeric animals were generated and utilized three months later as recipients of allogeneic HCT. The allogeneic [WT B6Ly5.2 → *Sdhaf1^-/-^* B6] chimeras demonstrated significantly shorter survival when compared with WT B6Ly5.2→ WT B6 recipients (Fig. 6j).

### In vitro and in vivo mechanism of T cell mediated reduction of SDHA in enterocytes

To investigate the mechanism of T cell-dependent reduction in SDHA, we first analyzed whether the reduction of SDHA in IECs was dependent on direct contact with T cells. The PCECs targets were incubated with T cells primed with syngeneic or allogeneic stimulators in a transwell assay. PCECs cultured with primed syngeneic T cells, as expected, did not show any increase in apoptosis regardless of contact. By contrast, the PCECs that were in direct contact with allo-primed T cells, showed greater apoptosis when compared to those separated by transwell membrane from the allo-primed T cells (Fig. 7a). Importantly when the PCEC targets were isolated and stained with SDHA from these transwell experiments after 12 hours of culture, only the PCECs that were in direct contact with allo-primed T cells showed reduction in SDHA when compared to those separated by transwell from allo-primed T cells or syn-primed T cells (Fig. 7b). These data demonstrated that reduction of SDHA in the IEC targets required direct contact by T cells primed against allo-antigen and that soluble mediators such as inflammatory cytokines were not the cause for reduction in SDHA levels in the IEC targets.

**Figure 7:**
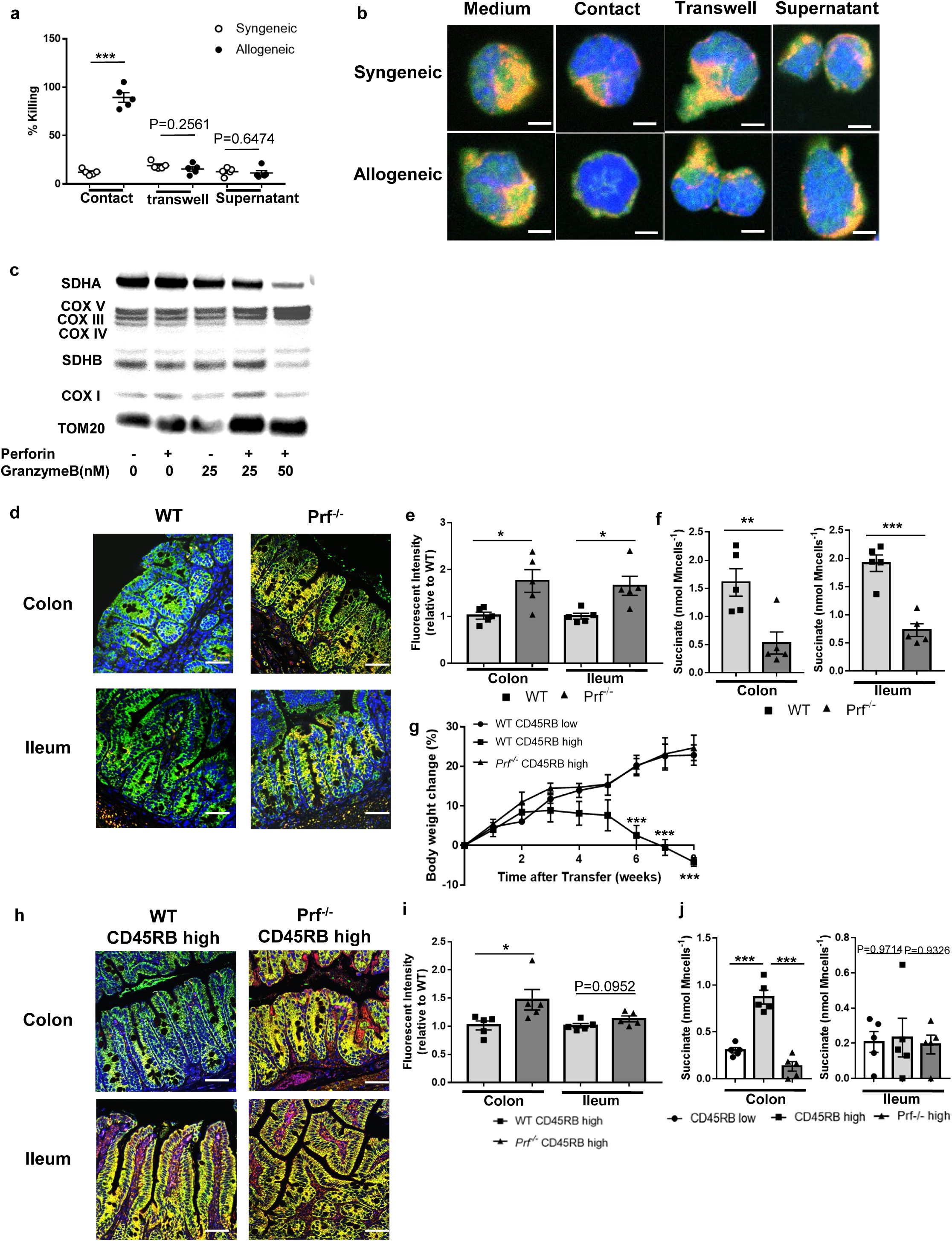
Granzyme B dependent abrogation of SDHA levels in IECs. (a-b) (**a**) Isolated splenic T cells from either B6 WT or BALB/c mice were cultured with irradiated splenocytes derived from B6 WT for 96 hours and CD8+ T cells were purified as effector cells. against PCECs was determined in direct or indirect co-culture with T-cells (using transwell membrane) or supernatant treatment from effector cell culture after 12 hour treatment (n=5 biological replicates and n=2 technical replicates). (**b**) Representative images of immunofluorescence staining of CD326+ PCECs from T cell treatment as shown in (**a**) (Complex IV=green, SDHA=red, DAPI=blue, scale bar= 5μm, n=5). (**c**) Immunoblot of SDHA, Complex V, IV, III, SDHB, I and TOM-20 in PCECs treated with perforin and Granzyme B for 4 hours. (d-f) BALB/c WT animals received 8 Gy total body irradiation and received 1×10^6^ T cells and 5×10^6^ TCD-BM cells from B6 WT or Prf-/-donors. (**d**) Representative images of immunofluorescence staining and (**e**) fluorescent intensity of colon and ileum from BALB/c mice that received B6 WT or Prf-/-donor cells 21days post allo-HCT (Complex IV=green, SDHA=red, DAPI=blue, scale bar= 50 μm, n=5). (**f**) Succinate levels in isolated IECs from colon and ileum of BALB/c mice received B6 WT or Prf-/-donor cells 21days post allo-HCT (n=5). (g-j) Naϊve T cells from B6 WT or Prf-/-mice or non-naϊve T-cells from B6 WT mice were transferred to RAG-1-/- mice. (**g**) Time course of body weight changes after T-cell transfer (n=5). (**h**) Representative images of immunofluorescence staining and (**i**) fluorescent intensity of colon and ileum 8 weeks after induction of colitis (scale bar= 50 μm, n=5). (**j**) Succinate levels in isolated IECs from colon and ileum of RAG-1-/- mice 8 weeks after T cell transfer (n=5). Representative plots and a graph summarizing the results of at least two independent experiments are shown. All statistical analysis by unpaired t-test (a, f), Mann-Whitney test (e, i) and one-way ANOVA analysis with Tukey post hoc test (g, j) (mean ± s.e.m.) were used to determine significance: *P < 0.05, **P < 0.01, ***P < 0.001.

We therefore next reasoned that T cell cytotoxic pathways might be the cause for reduction in SDHA. Previous reports have suggested that granzyme B (GzmB) degrades mitochondrial complex proteins in parasites^25^. Therefore, we next hypothesized that GzmB was required for reduction in SDHA. We cultured PCECs with increasing concentrations of recombinant perforin (PFN) and/or GzmB. The combination of both GzmB and PFN caused a decrease in SDHA levels in the PCECs when compared with control, PFN, or GzmB alone (Figure 7c). Furthermore, mitochondrial ROS production was increased when treated with both GzmB and PFN (Supplementary Fig. 7a). These data suggest that the requirement of perforin dependent intracellular transport of GzmB is required for reducing SDHA.

To further dissect the direct mechanisms GzmB mediated reduction of SDHA, we next hypothesized that GzmB directly cleaves SDHA once it gains access to the IEC cytoplasm following effects of perforin. SDHA, but not SDHB, demonstrates the putative GzmB recognition Ile-Glu-Asp↓Gly site (Supplementary Fig. 7b) ^26^. Mitochondria isolated from IECs were treated ex vivo with perforin and GzmB. Treatment with GzmB caused proteolysis of SDHA with a reduction in SDHA but an increase in SDHA cleaved fragments (supplementary Fig. 7c). Next, to rule out confounding effects from other cellular proteins, and to directly assess proteolysis of SDHA by GzmB, we utilized cell free experimental system. Recombinant SDHA was treated with GzmB and assessed for its full length and cleaved peptides. Recombinant SDHB was used as control. GzmB caused direct proteolysis of recombinant SDHA but not SDHB (Supplementary Fig. 7d) demonstrating that the mechanism for reduction of SDHA is because of its proteolysis by GzmB released from the T cells.

T cells utilize other cytotoxic pathways, including FasL, which could induce target cell apoptosis. Therefore to determine whether Fas-Fasl pathway could mediate similar enterocyte damage, we utilized B6 FasL deficient donor T cells with WT BM in the allo-HCT model as above. FasL deficient donor T cell caused similar reduction as WT T cells in allogeneic recipients (supplementary Fig. 7 f).

Taken together, these data reveal that T cells directly induce IEC damage and that GzmB/PFN play critical roles in abrogating SDHA levels in vitro and in vivo.

### Clinical correlation of loss of SDHA in humans

We next determined whether the loss of SDHA has any human, clinical relevance. To this end we obtained intestinal biopsy samples from 31 patients that underwent allogeneic HCT and received similar immunoprophylaxis and were clinically suspected for having lower GI GVHD. They all then underwent GI biopsy for diagnostic confirmation. None of these patients were on mycophenolate mofetil at the time of biopsy. In about half of these patients, biopsy confirmed the clinical diagnosis and in the other half, the biopsy was read as negative for GI GVHD. These 31 samples were stained and quantified for SDHA expression and analyzed in a manner blinded to the clinical and histopathological diagnosis. SDHA expression was dramatically reduced in the colonic biopsy samples from patients that were histopathologically confirmed as GI GVHD when compared with those who were pathologically considered not to have GI GVHD (Fig. 8a, Supplementary Fig. 8). The SDHA fluorescence intensity levels was statistically significant between GVHD and non-GVHD samples (Fig. 8b).

**Figure 8:**
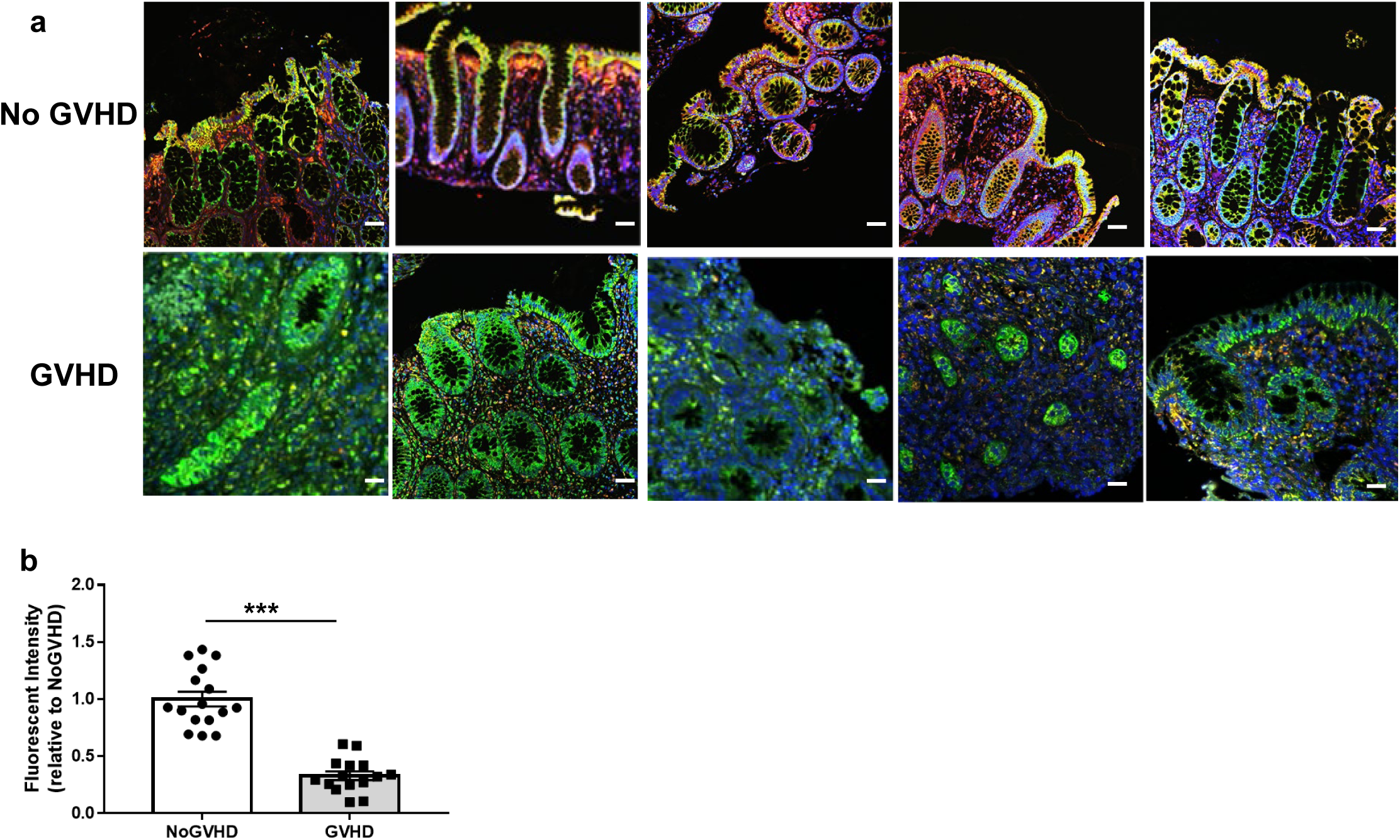
Clinical correlation of SDHA levels in IECs from human clinical samples. (a-b) Representative images of immunofluorescence staining of colonic biopsy samples from patients clinically suspected to have lower GI GVHD after allo-HCT (a, scale bar= 50 μm, GVHD: n=15, non-GVHD: n=16) and (**b**) Integrated fluorescent intensity of SDHA levels. All statistical analysis by unpaired t-test (**b**) (mean ± s.e.m.) were used to determine significance: ***P < 0.001.

## DISCUSSION

The metabolism of immune cells, such as T cells is dysregulated in immune diseases. T cells play a key role in alloimmune, autoimmne and iatrogenic colitis. The critical T cell and other immune cell dependent mechanisms between these various T cell mediated diseases such as GI GVHD, IBD and ICB are being increasingly recognized and understood. However, the tissue intrinsic mechanisms the immune targets, the IECs, remain uncharacterized. Importantly, whether the target cell intrinsic metabolic aberrations can regulate the overall disease severity of colitis remain unknown. Here we demonstrate a specific metabolic aberration in mitochondrial complex II of the IECs that is common to and exclusive for many T cell mediated diseases of the intestinal tract. We show that the IECs accumulate succinate as a consequence of reduction in their mitochondrial complex II component, SDHA. This reduction in SDHA caused an enhanced sensitivity of the IECs to T cell mediated cytotoxicity, thus serving as an IEC intrinsic metabolic checkpoint that regulated disease severity independent of direct effects of immune cells. The inhibition of SDHA in IECs altered their bioenergetics, decreased OXPHOS without a compensatory increase in glycolysis, which led to reduced O2 utilization, ATP generation and an enhanced ROS production. GzmB/PFN from the reactive T-cells worked together to lyse SDHA in IECs. PFN disrupt host cells to deliver the GzmB, which degrades SDHA and disrupts mitochondrial complex II and thus inhibits both TCA cycle and mitochondrial respiration of the IECs. Utilizing combination of metabolic, genetic, chemical loss and gain of function approaches we demonstrated that SDHA reduction in IECs is the critical target tissue intrinsic common pathway for T-cell mediated colitis, specifically in five models of GI GVHD, two (DSS and CD45RB^high^) models of autoimmune colitis, and a CTLA-4Ig ICB mediated colitis models. We further validated the observation in human samples of GI GVHD. These data identify tissue intrinsic SDHA as novel target to mitigate multiple T cell mediated intestinal immunopathologies^27^.

Succinate is a metabolic intermediate that is converted into fumarate by SDHA in the TCA cycle. Succinate plays an important role as an inflammatory signal, or as a source for reverse electron transport (RET) in the context of preserved SDH functions ^28^. In our study the impact on complex II from reduction of SDHA demonstrates that both TCA cycle and mitochondrial respiration are affected ^29^. Eukaryotes mainly produce energy as ATP via glycolysis in the cytoplasm or TCA cycle and ETC in mitochondria depending on cell status and oxygen levels. T cells (other immune cells as well), when activated show an increase in both glycolysis and OXPHOS^30 31^. By contrast, the intestinal immune targets, IECs show a loss of their major energy pathway, TCA cycle, without a compensatory change in glycolysis leading to a metabolic shutdown, which likely contributes to their greater susceptibility to apoptosis. At homeostasis, IECs efficiently utilize β-oxidation via SCFAs, especially butyrate, and SDHA serves as a part of TCA cycle enzyme and as a part of ETC ^15^. However, in the context of T cell colitis, not only did glycolysis levels show no changes, but flux studies demonstrated that neither glucose, nor odd chain carnitine nor glutamine supplementation failed to complement or compensate the energy production in IECs. These results show that IECs critically depend on OXPHOS and that neither enhanced glycolysis nor anaplerosis mitigated the defects from the reduction of SDHA. Energy insufficiency in IECs enhanced susceptible to T-cell mediated cytotoxicity was observed in vivo with both chemical inhibition and IEC specific genetic ablation of SDHA studies. Thus succinate accumulation in IECs caused by SDHA reduction, i.e defect in mitochondrial complex II, is distinct from the mitochondrial defects observed in other contexts such as ischemia reperfusion model, LPS activated macrophages or in other mitochondrial complex deficiencies ^21 22 32^. However, decreased NADH production by insufficient TCA cycle, a substrate for complex I reaction the starting point of ETC, could also secondarily contribute to further slowing down of the mitochondrial respiration and metabolic shutdown in the IECs caused by pathogenic T cells.

Acute GVHD, intriguingly, affects only the GI tract, liver and skin, the mechanisms for which remain unclear. Nonetheless, clinically, GI GVHD is the major cause for fatality from acute GVHD. Our data demonstrate that IECs when targeted by alloimmune cells across multiple models, regardless of type of allo-disparity and type of conditioning regimen, demonstrated reduction in SDHA with a concomitant increase in succinate. Importantly this reduction in SDHA was observed in human IECs only in those who had histopathologically confirmed GI GVHD. These data suggest that SDHA levels in IECs may serve as a marker for GI GVHD. Future, well designed human studies will evaluate its role as a potential diagnostic or prognostic biomarker for GI GVHD. Importantly, the reduction in SDHA was observed on in the IECs but not in the non-target organs of GVHD such as the kidneys demonstrating that specificity of T cell mediated cytotoxicity as a critical feature. This is further corroborated by lack of the effect on SDHA/succinate in the context of conditioning initiated damage such as from radiation induced colitis in the absence of alloreactive T cells. Intriguingly this specific mitochondrial abnormality was also not observed in other known GVHD target organs, namely liver and skin (data not shown), which are also caused by alloreactive T cells. This suggests that the dominant metabolic pathways and their alterations in IECs may be distinct from other GVHD target tissues. Future studies will need to systematically characterize the metabolic aberrations, if any, in these and all other organ cell types in the context of stress, inflammation, and T cell mediated cytotoxicity. It is also possible that instead of, or in addition to, the metabolic dysregulations, other critical survival pathways may also be distinct in the various GVHD target organ.

The specificity of mitochondrial complex II component, SDHA, dysregulation was also observed only in the context of T cell mediated autoimmune colitis and not in a chemically induced (DSS) colitis model wherein colitis is induced by both innate and adaptive immune cells^33^. Additionally, it was observed only when T cells were activated by checkpoint blockade in the context of low dose DSS resulting in colitis. This model reflects ICB mediated colitis, a major complication of the checkpoint blocker therapies against cancer^19, 20^. Importantly, inhibition of SDHA aggravated autoimmune and ICB mediated colitis. Thus the dysregulation of complex II of mitochondria in the IECs is a critical metabolic checkpoint for alloimmune, autoimmune and iatrogenic T cell mediated colitis.

These data also raise several interesting possibilities with potentially significant implications. For example, it is possible that T cell mediated autoimmune diseases afflicting other organs may share this common pathway. These data align with the notion that factors that promote tissue tolerance such as with growth and differentiation factor15 (GDF-15) in infectious diseases and extend the concept to non-infectious T cell mediated colitis ^37^. They demonstrate for the first time a tissue autonomous role for regulation of non-infectious immune-pathology supporting the concept that optimal therapy for immune mediated diseases will need approaches that promote immune tolerance and tissue resilience. The data thus suggest that mitochondrial SDHA in IECs may represent a novel target to mitigate the severity of GI GVHD, IBD and ICB colitis as an adjunct to T cell immunoregulatory approaches^27, 38–42^.

Furthermore, the converse, inhibiting SDHA in T cell targets may have therapeutic implications for enhancing T cell dependent immunotherapies. For example, it is possible that inhibiting SDHA in cancers where the tumor cells are predominantly dependent on OXPHOS may enhance their susceptibility to graft versus leukemia or ICB mediated cancer immunotherapy. Future studies will need to systematically explore these possibilities. Another intriguing possibility is that SDHA disruption in IECs led to reduced OXPHOS and ETC and caused decreased utilization of O_2_, potentially leading to increase in O_2_ levels in the intestinal lumen. This increase in O_2_ may provide a potential mechanistic explanation for changes in microbiome associated with GVHD and IBD. The change in O_2_ levels might lead to dysbiosis with loss of microbial diversity, decrease in commensals such as obligate anaerobes and shift towards to aerobes and pathobionts in the microbiome of patients with GVHD and IBD^43 44 45 46 47 48 49 50 51 52 53 54 55^.

In summary we identify a novel IEC intrinsic metabolic checkpoint that regulates their sensitivity to T cell mediated cytotoxicity and identify the component of mitochondrial complex II, SDHA, as a potential novel therapeutic target to regulate alloimmune, autoimmune and iatrogenic T cell mediated colitis.

## METHODS

### Mice

C57BL/6 (WT B6, H-2K^b^, CD45.2), B6 Ly5.2 (H-2K^b^, CD45.1), *B6.129S7-Rag1^tm1Mom^/J*(*Rag-1^-/-^*), *C57BL/6-Prf1^tm1Sdz^/J(Prf^-/-^), B6Smn.C3-Faslgld/J (FasL^-/-^), Sdhaf1^-/-^*, *B6.Cg-Tg(ACTFLPe)9 205Dym/J*, *B6.Cg-Tg(Vil-cre)1000Gum/J* mice and C3H.SW (H-2K^b^, CD229.1) mice were purchased from the Jackson Laboratory (Bar Harbor, ME, USA). BALB/c (H-2K^d^) and BDF1 (H-2K^b/d^) mice were purchased from Charles River Laboratories (Wilmington, MA, USA). C57BL/6N-*Sdha^tm2a(KOMP)Wts i^* mice were obtained from Knock Out Mouse Project (KOMP) repository, University of California, Davis (Davis, CA, USA) and bred to *ACTFLPe* mice to excise the *FRT-*flanked region (Supplementary Figure 6f). Then *Sdha^fl/fl^* mice were bred to *Vil1-Cre* mice to create *Vil1-Cre Sdha^fl/fl^* (*Sdha*^Δ^) mice. 8-12 weeks old female mice used for experiments. *All* mice were kept under specific pathogen-free (SPF) condi tions and cared for according to regulations reviewed and approved by the University of Michigan Committee on the Use and Care of Animals, which are based on the University of Michigan Laboratory Animal Medicine guidelines.

### Cell culture

C57BL/6 primary colonic epithelial cells (PCEC, C57-6047, Cell Biologics, Chicago, IL) were cultured in compete epithelial growth medium supplemented with insulin-transferrin-selenium, epi dermal growth factor, glutamine, antibiotics, antimycotics and fecal bovine serum (#M6621, Cell Biolog ics). Cells were routinely tested for mycoplasma contamination using MycoAlert (#LT07-318, Lonza, Switzerland). Cells were treated with Mouse active perforin (#APB317Mu01, Cloud-Clone Corp., Katy, TX) and recombinant mouse Granzyme B (#140-03, PeproTech, Rocky Hill, NJ) for 4 to 6 hours in indi cated concentrations following manufacturer’s recommendations.

### Generation of bone marrow (BM) chimeras

WT B6 and *Sdhaf1*^−/−^ animals were subjected to 1000 cGy total-body irradiation (TBI) from a ^137^Cs source on day −1 and then injected intravenously with 5 × 10^6^ BM cells from B6 Ly5.2 donor mice on day 0. Donor hematopoietic chimerism was confirmed using a CD45.2 monoclonal antibody 3 months after transplantation.

### Hematopoietic cell transplantation (HCT)

Splenic T cells from donors were enriched, and the BM was depleted of T cells by autoMACS (Miltenyi Biotec, Bergisch Gladbach, Germany) utilizing CD90.2 microbeads (#130-121-278, Miltenyi Biotec). WT B6, C3H.SW, *Sdhaf1*^-/-^, *Sdha*^Δ^ and BALB/c animals were used as recipients and received 1,000 cGy (WT B6, *Sdhaf1*^-/-^ and *Sdha*^Δ^) or 800 cGy (BALB/c) TBI on day −1, respectively, and 1 × 10^6^ (C3H.SW→ C57BL/B6 or C57BL/B6→BALB/c), 3× 10^6^ (BALB/c→C57BL/6, *Sdhaf1*^-/-^, *Sdha*^Δ^) CD90.2 T cells along with 5 × 10 T-cell-depleted BM→ (TCD-BM) cells from either syngeneic or allogeneic donors on day 0. [B6Ly5.2→ 6] and [B6 Ly5.2 → *Sdhaf1*^-/-^] animals received 900 cGy TBI on day −1 and were injected intravenously with 3 × 10^6^ CD90.2^+^ T cells and 5 × 10^6^ TCD-BM from either syngeneic B6 or BALB/c donors on day 0. For the chemotherapy conditioning model, B6 WT mice received busulfan (#B2635, 25 mg kg^-1^ from day −7 to −4; Sigma-Aldrich) and cyclophosphamide (#C7397, 100 mg kg^-1^ from day −3 to −2; Sigma-Aldrich) intraperitoneally, and 1 × 10^7^ (BALB/c→C57BL/6) CD90.2^+^ T cells along with 1 × 10^7^ TCD-BM cells from either syngeneic or allogeneic donors on day 0. For the no-irradiated model, unirradiated B6D2F1 mice were intravenously injected with 10 × 10^7^ splenocytes from B6D2F1 or B6 donors. Animals received vehicle or dimethyl malonate (DMM, #136441, 0.5g or 5g kg^-1^, Sigma-Aldrich, St. Louis, MO, USA), dimethyl itaconate (DI, #09533, 0.25g or 2.5g kg^-1^, Sigma-Aldrich) or atpenin A5 (#11898, 0.9µg or 9µg kg^-1^, Cayman Chemical, Ann Arbor, MI, USA) according to manufacturer’s instructions by flexible 20-gauge, 1.5-in. intra-gastric gavage needle every other day from day 0. The mice were randomly assigned to syngeneic, allogeneic or treatment groups in each experiment. No mice were excluded from analysis. No statistical methods were used to predetermine sample size. The investigators were not blinded to allocation during experiments and outcome assessment.

### Systemic and histopathological analysis of GVHD

**S**urvival after HCT was monitored daily and assessed the degree of clinical GVHD weekly, as described previously ^56^. Histopathological analysis of the liver, gastrointestinal (GI) tract, and lung, which are the primary GVHD target organs, was performed as described utilizing a semi-quantitative scoring system implemented in a blinded manner by a single pathologist (C.L.) ^57^. After scoring, the codes were broken, and the data compiled.

### Colitis models

For the DSS colitis model, animals were provided with drinking water containing 2.5-3% DSS (#0216011010, MP Biomedicals, Santa Ana, CA) for 7 days. Mice were injected with 100µg of anti–CTLA-4 mAb (#BE0164, clone 9D9, Bio X Cell, West Lebanon, NH) or isotype control (#BE0086, MCP-11, Bio X Cell) twice (3 and 1 day before DSS administration). For the T-cell transfer induced colitis model, isolated splenic T cells from WT B6 and *Prf^-/-^* mice were stained with DAPI (#422801, 1µM, Biolenged), APC-Cy7 anti-CD4 (#560246, GK1.5, 1:100, BD Biosciences, San Jose, CA), APC anti-CD25 (#101910, 3C7, 1:100, Biolegend), FITC anti-CD44 (#103006, IM7, 1:100, Biolegend) and PE anti-CD45RB (#103308, C363-16A, 1:100, Biolegend). CD4^+^CD25^-^CD44^-^ CD45RB^hi^ cells were sorted with the MoFlo Astrios cell sorter (Beckman Coulter, Indianapolis, IN) and intraperitoneally injected into *Rag-1^-/-^* recipients.

### Liquid chromatography mass spectrometry (LC/MS)

To quantitate tricarboxylic acid cycle metabolites in vivo, samples (colon, ileum and kidney) from mice 7 and 21days post HCT were harvested, homogenized, and snap-frozen in liquid N_2_. Tricarboxylic acid metabolites were extracted from tissue homogenates with a mixture of methanol, chloroform, and water (8:1:1) containing C13 isotope-labeled internal standards for citrate, succinate, fumarate, malate, alpha-ketoglutarate, lactate and pyruvate. Liquid chromatography-mass spectrometry (LC/MS) analysis was performed on an Agilent system consisting of a 1260 UPLC module coupled with a 6410 Triple Quadrupole (QQQ) mass spectrometer (Agilent Technologies, Santa Clara, CA). Data were processed using MassHunter Quantitative analysis version B.07.00. Metabolites were normalized to the nearest isotope-labeled internal standard and quantitated using a linear calibration curve ^58, 59^.The tissue levels were normalized by the protein concentration of the homogenized tissues.

### Acyl-carnitine quantitation

Tissues were subjected to targeted metabolomics analysis by LC/MS for determination of acyl-carnitines as previously described^60^. Briefly, samples were homogenized in 25mM phosphate buffer (pH 4.9) and extracted with cold 2:1:1 isopropanol: acetonitrile: methanol. Known amounts of isotope-labeled carnitines were used as internal standards and analyzed for LC/ESI/MS/MS analysis, an Agilent 6410 triple quadruple MS system equipped with an Agilent 1200 LC system and electrospray ionization (ESI) source was utilized and were detected in the multiple reaction monitoring (MRM) mode and relative peak areas were obtained.

### Metabolic flux analysis (MFA) assessing label incorporation into TCA cycle metabolite in vivo

Animals 7 and 21days post HCT were intragastrically gavaged with a bolus (2 g/kg) of either labeled ^13^C-glucose (#CLM-1396-1, Cambridge Isotopes, Tewksbury, MA) or non-labeled ^12^C-glucose after 9h fasting. The small and large intestinal epithelial cells were then isolated 4 hours later and prepared for analysis as above. The incorporation of ^13^C-glucose into the TCA cycle metabolites in the intestinal tissue was measured using LC/MS performed on Agilent 6520 high resolution Q-TOF (quadrupole-time of flight instrument) coupled with an Agilent 1200 HPLC system (Agilent Technologies, New Castle, DE), equipped with an electrospray source. The extract was subjected to hydrophilic interaction chromatography using Phenomenex Luna NH_2_ column (particle size 3 µm; 1 × 150mm) at a flow rate of 0.07 mL/min. Solvent A was 5 mM ammonium acetate with pH 9.9 and solvent B was acetonitrile. The column was equilibrated with 80% solvent B. The gradient was: 20-100% solvent A over 15 min; 100% solvent A over 5 min; 20% solvent B for 0.1 min; and 20% solvent A for 15.9 min. Liquid chromatography electrospray ionization (LC/ESI) MS in the negative mode was performed by the Q-TOF instrument using the following parameters: spray voltage 3000 V, drying gas flow 10 L/min, drying gas temperature 350°C, and nebulizer pressure 20 psi. Fragmentor voltage was 150 V in full scan mode. Mass range between *m/z* 100 to 1500 was scanned to obtain full scan mass spectra. Two reference masses at *m/z*121.050873 and *m/z* 922.009798 were used to obtain accurate mass measurement within 5 ppm. All chromatograms and corresponding spectra of TCA metabolites: citrate, succinate and malate and their corresponding ^13^C labeled counterparts were extracted and deconvoluted using MassHunter software (Agilent Technologies, New Castle, DE). Retention time consistency was manually rechecked and compared to authentic compounds that were injected under similar chromatographic conditions. For tissue extracts, metabolite concentrations were normalized to protein content, which was determined by the Bradford-Lowry method. For the flux analyses, peak areas of the labeled compounds were normalized to natural abundance of the label and represented as ratios to the total compound peak area.

^13^C-glucose or ^13^C-glutamine (#CLM-1822, Cambridge Isotopes) tracing in vitro was performed using glucose-free DMEM (#A1443001, Thermo Fisher Scientific, Waltham, MA) supplemented with 1mM sodium pyruvate (#11360070, Thermo Fisher Scientific), 1% HEPES (#15630080, Thermo Fisher Scientific), 2% BSA (#126575, Sigma-Aldrich), 10uM Y-27632 (#Y0503, Sigma-Aldrich) and either 17.5mM ^12^C or ^13^C-glucose, and either 2mM ^12^C or ^13^C-glutamine. Isolated intestinal epithelial cells (IECs) from naïve, syngeneic and allogeneic animals 7 and 21 days post HCT were cultured for 2 hours in ^12^C or ^13^C glucose/glutamine labeling media. Intracellular metabolite fractions were prepared from cells that were lysed with cold (−80°C) 80% methanol, then clarified by centrifugation. Metabolite pellets from intracellular fractions were normalized to the protein content of a parallel sample, and all samples were lyophilized via speed vac. Dried metabolite pellets from cells were re-suspended in 35 μ 50:50 MeOH: H_2_O mixture for metabolomics analysis. Agilent 1260 UHPLC combined with a 6520 Accurate-Mass Q-TOF LC/MS was utilized. Agilent MassHunter Workstation Software LC/MS Data Acquisition for 6200 series TOF/6500 series QTOF (B.06.01) was used for calibration and data acquisition. A Waters Acquity UPLC BEH amide column (2.1 x 100mm, 1.7μ phase (A) consisting of 20 mM NH_4_OAc in water pH 9.6, and mobile phase (B) consisting of acetonitrile. The following gradient was used: mobile phase (B) was held at 85% for 1 min, increased to 65% at 12 min, then to 40% at 15 min and held for 5 min before going to the initial condition and holding for 10 min. The column was cooled to 40°C and 3 μ MS with a flow rate of 0.2 mL/min. Calibration of TOF MS was achieved using the Agilent ESI Low Concentration Tuning Mix. Key parameters for both acquisition modes were: mass range 100-1200 da, Gas temp 350°C, Fragmentor 150 V, Skimmer 65 V, Drying Gas 10 L/min, Nebulizer at 20 psi, Vcap 3500 V and Ref Nebulizer at 20 psi. For the negative mode the reference ions were at 119.0363 and 980.01637 m/z whereas for positive acquisition mode, reference ions at 121.050873 and 959.9657 m/z. For data analysis, we used Agilent MassHunter Workstation Software Profinder B.08.00 with Batch Targeted Feature Extraction and Batch Isotopologue Extraction and Qualitative Analysis B.07.00.

Various parameter combinations, e.g., mass and RT tolerance, were used to find the best peaks and signals by manual inspection. Key parameters were: mass tolerance = 20 or 10 ppm and RT tolerance = 1 or 0.5 min. Isotopologue ion thresholds and the anchor ion height threshold were set to 250 counts and the threshold of the sum of ion heights to 500 counts. Coelution correlation threshold was set to 0.3^61^.

### Cell isolation

Luminal contents from dissected colon and ileum were flushed with CMF buffer; Ca^2+^/Mg^2+^ free HBSS (#14185052, Thermo Fisher Scientific) supplemented with 25mM sodium bicarbonate (#S6014, Sigma-Aldrich) and 2% FBS (#100-106, Gemini Bio Products, USA). Intestines were then minced into 5mm pieces, washed with CMF buffer four times, transferred to CMF with 5mM EDTA (#51201, Lonza, USA), and incubated at 37 °C for 40 minutes (shaking tubes every 10 minutes).

Supernatants containing IECs were then transferred through 100 µM cell filter followed by incubation on ice for 10 minutes to allow sedimentation. Supernatants were again transferred through a 75 µM cell filter. CD326^+^ IECs were next purified with APC-anti-CD326 (G8.8, #118214, 1:200, Biolegend) and anti-APC magnetic microbeads (#130-090-855, Miltenyi Biotec).

### Succinate dehydrogenase functional staining

Specimens were dissected from naïve, syngeneic and allogeneic WT B6 mice at 7 and 21 days post HCT. The tissues were rinsed in ice cold PBS and were then frozen in 2-methylbutane (#M32631, Sigma-Aldrich) and dry ice. Frozen sections (cut at 8µm thick) were directly placed in SDH solution; 48mM disodium succinate hexahydrate (#8201510100, Sigma-Aldrich), 0.75mM sodium azide (#S8032, Sigma-Aldrich), 0.5mM disodium EDTA (#324503, Sigma-Aldrich), 13mM sodium phosphate monobasic monohydrate (#S3522, Sigma-Aldrich), 87mM sodium phosphate dibasic heptahydrate (#S9390, Sigma-Aldrich), 998.9µM 1-methoxy-5-methylphenazinium methyl sulfate (M8640, Sigma-Aldrich), 1.5mM nitroblue tetrazolium (#484235, Sigma-Aldrich) at 37°C for 30 minutes and then the slides were rinsed in ddH_2_O. After mounting the slides, images were obtained using Olympus BX51M (Olympus, Japan) and the SDH activity was quantified using Image J (National Institutes of Health, Bethesda, MA, USA).

### Mitochondria Isolation

Isolated IECs from syngeneic and allogeneic recipients in mitochondria isolation buffer (MIB) were disrupted using a glass homogenizer with 2–3 strokes. MIB was composed of 70mM sucrose (#S0389, Sigma-Aldrich), 210mM mannitol (#M9546, Sigma-Aldrich), 5mM HEPES, 1mM EGTA (#E0396, Sigma-Aldrich), 0.5% BSA, pH 7.2 and protease inhibitor cocktail (#78429, Sigma-Aldrich). Homogenate was centrifuged at 800 g for 10 min at 4°C. Following centrifugation, fat/lipid was carefully aspirated, and the remaining supernatant was decanted through 2 layers of cheesecloth to a separate tube and centrifuged twice at 8000g for 15 min at 4°C. After removal of the light mitochondrial layer, the pellet was resuspended in MIB and centrifuged at 8500g for 10min. The final pellet was resuspended in a minimal volume of mitochondrial assay solution (MAS) and kept on ice. MAS comprises 70mM sucrose, 220mM mannitol, 10mM potassium monobasic phosphate (#P5655, Sigma-Aldrich), 5mM magnesium chloride (#M8266, Sigma-Aldrich), 2mM HEPES, 1mM EGTA, 0.2% BSA, and protease inhibitor cocktail at the pH of 7.2. Total protein (mg/ml) was determined via Pierce BCA Protein Assay (#23225, Thermo Scientific).

### Immunoblot analysis

Isolated mitochondria or IECs were lysed in RIPA buffer (#89901, Thermo Scientific). Equal amounts of proteins were loaded on 4-12% SDS-PAGE gel (#NP0321, Thermo Scientific), electrophoresed and subsequently transferred to a PVDF membrane (#ISEQ85R, Sigma-Aldrich) using a Bio-Rad semi-dry transfer cell (20 V, 1 h). Blots were incubated with anti-SDHA (#ab14715, 2E3GC12FB2AE2, 0.1μg/ml, Abcam, Cambridge, UK), Total OXPHOS rodent antibody cocktail (#ab110413, 1:250, Abcam), Tom20 (#42406, D8T4N, 1:1000, Cell Signaling Technology, Danvers, MA) and anti-β actin (#8226, mAbcam8226, 1:3000, Abcam) primary antibodies overnight at 4°C. Incubation with secondary anti-rabbit-HRP (#7074S, Cell Signaling Technology), anti-mouse-HRP (#sc-516102, Santa Cruz Biotechnology, Dallas, TX) was performed at room temperature for 1 hour. Bound antibody was detected using Super Signal ECL substrate (Thermo Scientific) and quantitated using ChemiDoc MP Imaging system (BioRad, Hercules, CA). Densitometric analysis was performed using Image J.

### SDHA cleavage assay

75 ug Isolated mitochondria in 25 ul EDB buffer (10mM HEPES, 50mM NaCl, 2mM MgCl_2_, 5mM EGTA and 1mM DTT) were incubated with 75 ng mouse Granzyme B at 37 degrees and then separated by SDS-PAGE ^62^. Mouse recombinant SDHA protein with His Tag (LS-G26030, LSBio, Seattle, WA) and mouse recombinant SDHB protein with His Tag (LS-G26026, LSBio) in EDB buffer (25ug in 8.3ul) were incubated with 25ng mouse Granzyme B at 37 degrees for 90 minutes.

### BN-PAGE

Isolated mitochondria was lysed in NativePAGE sample buffer (#BN2008, Thermo Scientific) containing 2% digitonin. 50ug of total protein was loaded on 3-12% NativePAGE gel (#BN1001, Thermo Scientific). Gels were stained by Colloidal Blue Staining Kit (#LC6025, Thermo Scientific). For the second dimension, SDS-PAGE, gel lanes from BN gels were placed onto top of a 4-12% NuPage 2D well (#NP0326, Thermo Scientific). After separation and blotting, membranes were probed with Total OXPHOS blue native antibody cocktail (#ab110412, 1:250, Abcam).

### Fluorescent immunohistochemistry staining

Sections were prepared as mentioned in the SDH functional staining method. The sections were fixed in 4% paraformaldehyde (#P6148, Sigma-Aldrich) for 30 minutes. Then, the sections were washed 3 times in PBS and permeabilized in PBS containing 0.3% Triton-X100 (#X100, Sigma-Aldrich) for 2 hours. Permeabilized sections were blocked for 2 hours (10% normal goat serum in PBS containing 0.15% Triton X-100) and incubated with primary antibodies overnight at 4°C. The following primary antibodies were used; anti-SDHA (#ab14715, 2E3GC12FB2AE2, 1:100, Abcam), COX IV (#ab16056, 20E8C12, 1:200, Abcam), Granzyme B (#NB100-684, 1:100, Novus Biologicals, Centennial, CO) overnight at 4. Incubation with secondary Cy3 anti-mouse-IgG (#405309, Poly4053, 1:250, BioLegend, San Diego, CA), Cy3 anti-rabbit-IgG (#406402, Poly4064, 1:250, BioLegend) and Dy-Light488 anti-rabbit-IgG (#406404, Poly4064, 1:250, BioLegend) was performed at room temperature for 1 hour. Slides were mounted with coverslips using Prolong Gold with DAPI (#P36931, Thermo Scientific). For human samples, formalin fixed paraffin embedded specimens from patients were cut to a thickness of 5 µm. Heat-induced antigen retrieval was performed with 10 mM sodium citrate buffer followed by staining with primary antibodies. Slides were imaged using a Nikon A1confocal microscope (Nikon, Melville, NY). SDH levels were quantified and measured in intensity from Cy3 positive epithelial cells using Image J. For primary mice colonic epithelial cells, cells were spun down and put on coverslips. After 10 minutes of incubation at 37°C, cells were fixed with 3% paraformaldehyde for 20minutes, quenched with 100mM Glycine (#G8998, Sigma-Aldrich) 3 times for 3 minutes and permeabilized in PBS containing 0.3% Triton-X100 for 1 minute. Then, the cells were blocked, stained with primary and secondary antibodies, and mounted.

### Transmission Electron Microscopy and immunogold immunohistochemistry

Colon and ileum were harvested, diced into 1mm cubes and fixed with 4% formaldehyde and 2.5% glutaraldehyde (#G5882, Sigma-Aldrich) in 0.1 M phosphate buffer (PB, pH 7.4) overnight. Next, the tissues were immersed in 0.2 M sucrose and 0.1 M PB, and then post-fixed in 1% osmium tetroxide (#75632, Sigma-Aldrich) in 0.1 M PB. Then, the tissue was dehydrated in ascending concentrations of ethanol, treated with propylene oxide, and embedded in Epon epoxy resin. Semi-thin sections were stained with toluidine blue for tissue identification. Regions of interest were selected, cut into ultra-thin section (70 nm thick) mounted on copper grids, and then stained with uranyl acetate and lead citrate. For immunogold immunohistochemistry, sections were blocked with blocking solution for goat gold conjugates (#905.002, AURION, Netherland) for 2 hours and then incubated with anti-SDHA antibody (1:10) overnight at 4°C. After rinsing, sections were incubated with 10nm gold particle conjugated anti-mouse IgG (H&L) (#810.022, 1:50, AURION) for 2 hours at room temperature and then fixed with 2.5% glutaraldehyde for 10 min before staining with 4% uranyl acetate for 15 minutes. The samples were examined using a JEM-1400 electron microscope (JEOL, Peabody, MA) at 80 kV. Images were recorded digitally using a XR401 camera system operated with AMT software (Advanced Microscopy Techniques Corp., Danvers, MA). Gold particles in the mitochondria were quantified in a blinded manner by L.L. After quantification, the codes were broken, and the data were compiled.

### MtDNA Copy Number Analysis

Total DNA was extracted from isolated IECs using the DNeasy Blood & Tissue Kit (#69504, QIAGEN, Germantown, MD). PowerUP SYBR green master mix (#A25742, Applied Biosystems, Foster City, CA) and the following primers were used; 5’- CAAACACTTATTACAACCCAAGAACA-3’ and 5’- TCATATTATGGCTATGGGTCAGG-3’ (*ND1; NADH:ubiquinone oxidoreductase core subunit 1*); 5’- AATCTACCATCCTCCGTGAAACC-3’ and 5’- TCAGTTTAGCTACCCCCAAGTTTAA-3’ (*D-loop1; displacement loop1*); 5’- CCCTTCCCCATTTGGTCT-3’ and 5’- TGGTTTCACGGAGGATGG-3’ (*D-loop2*) ^63^; 5’- TCCTCCGTGAAACCAACAA-3’ and 5’-AGCGAGAAGAGGGGCATA-3’ (*D-loop3*); 5’- GCTTTCCACTTCATCTTACCATTTA-3’ and 5’-TGTTGGGTTGTTTGATCCTG-3’ (*CytB: cytochrome B*); 5’-CACTGCCTGCCCAGTGA-3’ and 5’-ATACCGCGGCCGTTAAA-3’ (*16S*); 5’- AACGGATCCACAGCCGTA-3’ and 5’-AGTCCTCGGGCCATGATT-3’ (*ND4*); and 5’- GCTTAATTTGACTCAACACGGGA-3’ and 5’-AGCTATCAATCTGTCAATCCTGTC-3’ (*18S)*. All primers were verified for the production of a single specific PCR product via melting curve analysis.

### Succinate quantification

Isolated IECs were collected, processed and analysed using the Succinate Assay Kit (#ab204718, Abcam) according to the manufacturer’s protocol.

### Seahorse analysis

Analysis was performed on IECs after HCT. Cells were washed three times in complete seahorse XF assay medium (#103335-100, Aglient technologies, Santa Clara, CA) with 17.5 mM glucose, 1 mM sodium pyruvate, 2 mM glutamine, 2 %BSA 10uM Y-27632 and 1% penicillin- streptomycin (#516106, Sigma-Aldrich) adjusted to pH 7.4. Cells were plated at 8 × 10^4^ cells per well in a Seahorse assay plate, pretreated with matrigel (#354262, Corning, Corning, NY). Cells were equilibrated to 37 °C for 30 min before assay. Respiration profile was assessed in 96XF instrument with Mitostress assay as indicated upon cell treatment with oligomycin (#75351, 2µM, Sigma-Aldrich), FCCP (#C2990, 1µM, Sigma-Aldrich), rotenone (#R8875, 0.5μ (#A8674, 0.5μM, Sigma-Aldrich) ^14^.

### ATP detection assay

Isolated mitochondria from IECs were incubated in MAS supplemented with 5mM ADP (#A2754, Sigma-Aldrich), 10mM succinic acid (#S3674, Sigma-Aldrich) and 2µM rotenone on white walled 96- well plates (#3917, Corning) at 8 × 10^4^ cells per well in triplicate for 2 hours. The ATP level was measured using the Cell Titer Glo 2.0 luminescence assay (#G9243, Promega, Madison, WI). Luminescence was measured for 500ms using a SpectraMax M3 Microplate reader (Molecular Devices, San Jose, CA).

### Complex I enzyme activity assay

Isolated mitochondria from IECs were loaded on 96-well plates at 5µg per well in triplicate. Enzyme activity was measured using the colorimetric Complex I Enzyme Activity Assay Kit (#ab109721, Abcam) following manufacturer’s instructions.

### NAD^+^/Nicotinamide Adenine Dinucleotide Diaphorase Assay

Total intracellular NAD NAD and NADH.

### ^51^Cr release assay

Splenic T cells from WT-B6 and BALB/c animals and irradiated (30Gy) red blood cell lysed splenocytes from WT-B6 mice were co-cultured in 5:2 ratio for 96 hours. Splenic CD8^+^ T cells were isolated from above co-cultured cells, purified with a CD8^+^ T-cell isolation kit (#130-117- 044, Miltenyi Biotec) and used as effector cells. PCECs were pre-treated with 5mM DMM, 0.5mM DI, 10nM atpenin A5 or vehicle for 3 hours. 2 × 10^6^ treated cells were incubated with 2 MBq of Na_2_ CrO_4_ (#NEZ030001MC, PerkinElmer, Waltham, MA) for 2 hours at 37 °C in a 5% CO_2_ atmosphere and were used as target cells. After washing, 4 × 10^3^ labeled targets were resuspended and added to triplicate wells at varying effector-to-target ratios and then incubated for 4 h. Maximal and minimum release was determined by the addition of Triton-X or media alone to targets, respectively.

For the transwell assays, 2 × 10^5^ labeled targets were placed on the bottom chamber and effector cells were on the bottom or on the 0.4uM transwell membrane for 12 hours at 37 °C in 5% CO_2_. Cells were incubated in fresh cell culture medium or conditioined from supernatant from effector cells. After incubation, supernatants were transferred to a Luma plate (#600633, PerkinElmer) and ^51^Cr activity was determined using Top Count NXT (Hewlett Packard, Palo Alto, CA).

### Flow cytometry

Cells were re-suspended in 2% BSA in PBS and stained with the following antibodies and reagents; FITC-anti-CD326 (G8.8, #118208, 1:200), APC-anti-CD326 (G8.8, #118214, 1:200), DAPI, CellROX Deep Red (#C10491, 500nM, Thermo Scientific), MitoSOX (#M36008, 5µM, Thermo Scientific), JC-1 (#M34152, 2µM, Thermo Scientific), 7-AAD (#420404, 1:100, BioLegend), Annexin V (#640947, 1:20, Biolegend) anti-CD16/CD32 antibody (BD Biosciences, 2.4G2) according to manufacturer’s protocol. Cells were run on Attune NxT flow cytometer (Thermo Scientific) and analyzed using FlowJo v10.2 (FlowJo LLC, Ashland, OR).

### RNA isolation and RT-PCR

Total RNA from single-cell suspensions was isolated using the RNeasy Kit (#74104, QIAGEN, Hilden, Germany) and reverse transcribed into cDNA using the High Capacity cDNA Reverse Transcription Kit (#4374966, Applied Biosystems). The following primers and PowerUP SYBR green polymerase were used to detect the following transcripts: 5′ ACATGCAGAAGTCGATGCAG-3′ and 5′ CATTCCCCTGTCGAATGTCT-3′ (Sdha); and 5′ -CTTCCCATTCTCGGCCTTG-3′ (Gapdh). All reactions were performed according to manufacturer’s instructions. All primers were verified for the production of a single specific PCR product viamelting curve analysis.

### Human Subjects and Data

Colonic biopsy samples embedded in paraffin with deidentified and blinded patient data diagnosed as GVHD or non-GVHD from the University of Michigan Department of Pathology was provided without any information. All protocols and procedures for the human studies were approved by the university Institutional Review Board.

### Statistical analysis

Experiments were conducted with technical and biological replicates at an appropriate sample size, as estimated by our prior experience. No statistical methods were used to predetermine sample size. No methods of randomization and no blinding were applied. All data were replicated independently at least once as indicated in the figure legends, and all attempts to reproduce experimental data were successful. Bars and error bars represent the mean and s.e.m., respectively. All statistical analysis was performed using Graph Pad Prism 7 (Graph Pad Software Inc, San Diego, CA). P values <0.05 were considered as significant: *P<0.05, **P<0.01, ***P<0.001, ****P<0.0001; P values >0.05 were considered as non-significant. All sample sizes and statistical tests used are detailed in each figure legend.

## Acknowledgments

This work was supported by the US National Institutes of Health grants HL090775, CA173878, CA203542 (P.R.), K08HL130944 (A.V.M), JSPS Postdoctoral Fellowships for Research Abroad (H.F.) and The YASUDA Medical Foundation Grants for Research Abroad (H.F.). We acknowledge use of the Microscopy & Image-analysis Laboratory (MIL) of the University of Michigan’s Biomedical Research Core Facilities for the preparation of samples and images. Support for the MIL core is provided by the University of Michigan Rogel Cancer Center (NIH grant CA46592).

## Author contributions

H.F. and P.R. conceived and designed this study. H.F. and P.R. planned and guided the research and wrote the manuscript. H.F. I.K., A.V.M., A.P., H.M., T.T., I.H., S.J.W., S.K., A.T., D.S., K.O.W., Y.S., and J.B. performed experiments. I.K., A.V.M., H.J.L. and J.B.analyzed LC/MS and MFA data. D.P. performed experiments, analyzed human data and edited the paper. C.L. performed experiments and histopathological analysis. H.F., T.S. and P.R. generated the *SDHA* floxed mice. P.S., T.S., C.A.L., S.P., A.R. and R.P. supervised the work carried out in this study.

## Conflict of interest statement

The authors have no conflict of interest.

## Supplementary Figure Legends

**Supplementary Figure 1: Analysis of ^13^C incorporation from ^13^C glutamine treatment in IECs post allo-HCT.** B6 WT animals received 10 Gy total body irradiation and received 3×10^6^ T cells and 5×10^6^ TCD-BM cells from either syngeneic B6 WT or allogeneic BALB/c donors. Abundance from uniformly ^13^C-glutamine of cis-aconitate, -ketoglutarate, succinate, and malate after 4αh incubation in syngeneic or allogeneic isolated IECs 7 and 21days post HCT (D7 Syn: n=4, Allo: n=5, D21 Syn: n=5, Allo: n=7). All statistical analysis for abundance by unpaired t-test (mean ± s.e.m.): \**P* < 0.05, \*\**P* < 0.001.

**Supplementary Figure 2: Specific reduction of SDHA activity in colon and ileum post allo-HCT.** B6 WT animals received 10 Gy total body irradiation and received 3×10^6^ T cells along with 5×10^6^ TCD- BM cells from either syngeneic B6 WT or allogeneic BALB/c donors. (**a**) Representative images of SDH enzyme activity staining in GVHD target tissues (colon, ileum, liver and skin) and non-target tissues (heart, pancreas and kidney) from naive animals or recipients 21days post HCT (Scale bar: yellow 500µm, black 200 µm). (**b**) Integrated intensity of SDH enzyme activity staining from colon and ileum 7days post HCT (n=5). Representative plots and a graph summarizing the results of at least two independent experiments are shown. All statistical analysis by Mann-Whitney test (**b**) (mean ± s.e.m.): \**P* < 0.05, \*\**P* < 0.01.

**Supplementary Figure 3: Morphological mitochondrial changes but not numbers in IECs post allo-HCT.** B6 WT animals received 10 Gy total body irradiation followed by 3×10^6^ T cells along with 5×10^6^ TCD-BM cells from either syngeneic B6 WT or allogeneic BALB/c donors. (**a**) Immunoblot protein density quantification of SDHA and SDHB in mitochondria from IECs 7days post HCT (n=4). (**b**) Representative images of immunofluorescence staining of kidney and liver from recipients 21days post HCT (Complex IV=green, SDHA=red, DAPI=blue, scale bar= 50 µm, n=5). (**c**) Numbers of gold particles per mitochondrial from colon and ileum of naïve animals or recipients 7days post HCT in transmission electron microscopy images with immune-gold staining of SDHA (total 50 mitochondria from 3 samples). (**d**) Representative images of transmission electron microscopy in mitochondria of colon from naive or recipients 21days post HCT (left; scale bar 200nm). Arrow indicates normal cristae and arrowhead indicates abnormal cristae (n=3). (**e**) Mitochondria DNA relative copy numbers of colon and ileum from syngeneic and allogeneic recipients 7 and 21days post HCT (n=5). Representative plots and a graph summarizing the results of at least two independent experiments are shown. All statistical analysis by Mann-Whitney test (**a, e**) or one-way ANOVA analysis with Tukey post hoc test (**c**) (mean ± s.e.m.): \*\*\*\**P* < 0.0001.

**Supplementary Figure 4: Conditioning regimen does not affect SDHA expression in IECs.** (**a-b**) B6 WT mice received 10 Gy total body irradiation followed by 1×10^6^ T cells along with 5×10^6^ TCD- BM cells from either syngeneic B6 or allogeneic mHA-micmatched C3H.SW donors (n=5). (**a**) Representative images of immunofluorescence staining and (**b**) fluorescent intensity of SDHA in colon and ileum from recipients 28days post HCT (scale bar=50µm) are shown. (**c-d**) B6 WT mice received chemotherapy and received 1×10^7^ T cells along with 1×10^7^ TCD-BM cells from either syngeneic B6 WT or allogeneic BALB/c donors (n=5). (**c**) Representative images of immunofluorescence staining and (**d**) fluorescent intensity of SDHA in colon and ileum from recipients 21days post HCT (scale bar=50µm). (**e-g**) B6 WT mice were treated with 3% DSS or vehicle in drinking water for 7 days (n=5). (**e**) Representative images of immunofluorescence staining and (**f**) fluorescent intensity of SDHA in colon 12days after DSS treatment (scale bar=50µm). (**g**) Succinate levels in isolated IECs from colon and ileum of mice treated with 3% DSS at day12. (**h**) B6 WT mice receiving isotype control IgG or anti- CTLA-4 antibody were treated with 3% DSS in drinking water for 7 days. Time course of body weight changes after DSS administration. Representative plots and a graph summarizing the results of at least two independent experiments are shown. All statistical analysis by Mann-Whitney test (**b, d, f**) or unpaired t-test (**g, h**) (mean ± s.e.m.): \**P* < 0.05, \*\**P* < 0.01.

**Supplementary Figure 5: SDH inhibition causes ROS accumulation in IECs.** (**a-e)** Primary colonic epithelial cells (PCECs) were treated with DMSO, malonate or itaconate for 6 hours (n=5). (**a-b**) Cytoplasmic ROS measured by CellROX staining. (**a**) Representative images and (**b**) CellROX positive cells are shown. (**c-d**) Mitochondria ROS measured by MitoSOX staining. (**c**) Representative images and (**d**) Cell SOX positive cells are shown. (**e**) PCECs were treated with DMSO or atpeninA5 for 4hours. Apoptosis (left), CellROX (middle) and MitoSOX (right) levels were determined (n=5). (**f-h**) B6 WT animals received 10 Gy total body irradiation followed by 3×10^6^ T cells along with 5×10^6^ TCD-BM cells from either syngeneic B6 WT or allogeneic BALB/c donors (n=5). (**f**) CellROX and MitoSOX levels in isolated colon and ileum IECs from syngeneic and allogeneic recipients 7 and 21days post HCT were analyzed. (**g**) Representative plots and (**h**) levels of mitochondrial membrane potential (ψm) in isolated colon and ileum IECs from syngeneic and allogeneic recipients 7 and 21days post HCT were analyzed. Representative plots and a graph summarizing the results of at least two independent experiments are shown. All statistical analysis by one-way ANOVA analysis with Tukey post hoc test (**b, d, e**) or unpaired t-test (**f, h**) (mean ± s.e.m.): \**P* < 0.05, \*\**P* < 0.01, \*\*\**P* < 0.001.

**Supplementary** figure 6: **Chemical inhibition and genetic ablation of SDHA in IECs regulate the severity of GVHD.** (**a**) Representative images of SDH enzyme staining of colon from naive B6 WT 12 hours after vehicle, malonate (5g kg^-1^) or itaconate (2.5g kg^-1^) treatment. (Scale bar 100µm, n=2). (**b-d**) B6 WT or *Sdhaf1^-/-^* animals received 10 Gy total body irradiation followed by 3×10^6^ T cells along with 5×10^6^ TCD-BM cells from either syngeneic B6 WT or allogeneic BALB/c donors. (**b**) Survival of B6 WT recipients treated with vehicle or atpeninA5 (9µg kg^-1^) every other day post HCT (n=8). (**c**) B6 WT recipients treated with vehicle, malonate (5g kg^-1^) or itaconate (2.5g kg^-1^) every other day post HCT. Pathological GVHD scores in liver, skin and lung 7days post HCT (n=6). (**d**) Pathological GVHD scores in liver, skin and lung 7days post HCT from B6 WT and *Sdhaf1^-/-^* recipients (n=6). (**e**) B6 WT and *Sdhaf1^-/-^* animals were treated with 2.5 % DSS in drinking water for 7 days. Time course of body weight changes after DSS administration (n=6). (**f**) Scheme illustrating the strategy used to generate *Sdha* floxed and excised alleles in IECs. (**g**) Expression of *Sdha* mRNA in CD326^+^ isolated IECs from six- week-old *Sdha*^Δ^ relative to *Sdha* mice (*n* = 4). *Sdha* expression was normalized to *GAPDH* expression. (**h**) Immunoblot of SDHA and TOM-20 in mitochondria of IECs from six-week-old *Sdha*^Δ^ and *Sdha* mice. (**i**) Representative images of immunofluorescence staining of colon, ileum, liver, skin and heart from six-week-old *Sdha*^Δ^ and *Sdha* mice (Complex IV=green, SDHA=red, DAPI=blue, scale bar=50µm, n=4). Representative plots and a graph summarizing the results of at least two independent experiments are shown. All statistical analysis by log-rank test (**b**), Mann-Whitney test (**d, e, g**) or Kruskal-Wallis analysis with Dunn’s post hoc test (**b**) (mean ± s.e.m.): \**P* < 0.05, \*\**P* < 0.01, \*\*\**P* < 0.001.

**Supplementary Figure 7: Granzyme B cleaves SDHA in mitochondria from IECs.** (**a**) PCECs treated with perforin and Granzyme B for 4hours were analyzed for CellROX and MitoSOX levels (n=4). (**b**) Amino acid sequences of SDHA and SDHB. A possible target by Granzyme B was shown in gray. (**c**) Isolated mitochondria were incubated with Granzyme B in indicated time (ON; overnight). Immunoblot image of full length or cleaved SDHA and cleaved/ full length SDHA ratio were shown (n=3, each). (**d**) His-Tagged recombinant SDHA and SDHB proteins were treated with Granzyme B for 90 minutes. Immunoblot image and ratio of cleaved/full length SDHA/ SDHB were shown (n=3, each). (**e**) BALB/c WT animals received 8 Gy total body irradiation followed by 1×10^6^ T cells along with 5×10^6^ TCD-BM cells from B6 WT or *Prf^-/-^*donors. Succinate levels in isolated IECs from colon and ileum of BALB/c mice that received B6 WT or *Prf^-/-^*donor cells 7days post allo-HCT (n=5). (**f**) BALB/c WT animals received 8 Gy total body irradiation followed by 1×10^6^ T cells along with 5×10^6^ TCD-BM cells from B6 WT or *FasL^-/-^*donors. Representative images of immunofluorescence staining and fluorescent intensity of colon from BALB/c mice that received B6 WT or *FasL^-/-^*donor cells 21days post allo-HCT (Complex IV=green, SDHA=red, DAPI=blue, scale bar= 50 µm, n=5). (**g-j**) Naïve T cells from B6 WT or *Prf^-/-^*mice or non-naïve T-cells from B6 WT mice were transferred to RAG-1^-/-^ mice (n=5). (**g**) Representative images of immunofluorescence staining of colon 8weeks after induction of colitis (Granzyme B=red, DAPI=blue, scale bar= 50 µm) and (**h**) Granzyme B positive cell numbers in colon. (**i**) CellROX and (**j**) MitoSOX levels were determined in colonic IECs 8weeks after induction of colitis (n=5). All statistical analysis by one-way ANOVA analysis with Tukey post hoc test (**a, d, e, f, g**) or unpaired t-test (**b, h**) (mean ± s.e.m.): \**P* < 0.05, \*\**P* < 0.01, \*\*\**P* < 0.001.

**Supplementary Figure 8: SDH expression in IECs from GVHD and non-GVHD patients.** Other representative images of immunofluorescence staining of colonic biopsy samples from patients suspected as having clinical GVHD after HCT not shown in Figure 8 (scale bar= 50 µm).

## References

1. Strober, W., Fuss, I. & Mannon, P. The fundamental basis of inflammatory bowel disease. The Journal of clinical investigation 117, 514–521, doi:10.1172/jci30587 (2007).

2. Yilmaz, B. et al. Microbial network disturbances in relapsing refractory Crohn’s disease. Nature medicine 25, 323–336, doi:10.1038/s41591-018-0308-z (2019).

3. Cleynen, I. et al. Inherited determinants of Crohn’s disease and ulcerative colitis phenotypes: a genetic association study. *Lancet (London*, England) 387, 156–167, doi:10.1016/s0140-6736(15)00465-1 (2016).

4. Jostins, L. et al. Host-microbe interactions have shaped the genetic architecture of inflammatory bowel disease. Nature 491, 119–124, doi:10.1038/nature11582 (2012).

5. Martins, F. et al. Adverse effects of immune-checkpoint inhibitors: epidemiology, management and surveillance. Nature reviews. Clinical oncology, doi:10.1038/s41571-019-0218-0 (2019).

6. Ferrara, J. L., Levine, J. E., Reddy, P. & Holler, E. Graft-versus-host disease. Lancet (London, England) 373, 1550–1561, doi:10.1016/s0140-6736(09)60237-3 (2009).

7. Neurath, M. F. Targeting immune cell circuits and trafficking in inflammatory bowel disease. Nature immunology, doi:10.1038/s41590-019-0415-0 (2019).

8. Zeiser, R. & Blazar, B. R. Acute Graft-versus-Host Disease - Biologic Process, Prevention, and Therapy. The New England journal of medicine 377, 2167–2179, doi:10.1056/NEJMra1609337 (2017).

9. O’Neill, L. A. & Pearce, E. J. Immunometabolism governs dendritic cell and macrophage function. The Journal of experimental medicine 213, 15–23, doi:10.1084/jem.20151570 (2016).

10. Everts, B. et al. TLR-driven early glycolytic reprogramming via the kinases TBK1-IKKvarepsilon supports the anabolic demands of dendritic cell activation. Nature immunology 15, 323–332, doi:10.1038/ni.2833 (2014).

11. Buck, M. D. et al. Mitochondrial Dynamics Controls T Cell Fate through Metabolic Programming. Cell 166, 63–76, doi:10.1016/j.cell.2016.05.035 (2016).

12. Albenberg, L. et al. Correlation between intraluminal oxygen gradient and radial partitioning of intestinal microbiota. Gastroenterology 147, 1055–1063.e1058, doi:10.1053/j.gastro.2014.07.020 (2014).

13. He, G. et al. Noninvasive measurement of anatomic structure and intraluminal oxygenation in the gastrointestinal tract of living mice with spatial and spectral EPR imaging. Proceedings of the National Academy of Sciences of the United States of America 96, 4586–4591, doi:10.1073/pnas.96.8.4586 (1999).

14. Fan, Y. Y. et al. A bioassay to measure energy metabolism in mouse colonic crypts, organoids, and sorted stem cells. Am J Physiol Gastrointest Liver Physiol 309, G1–9, doi:10.1152/ajpgi.00052.2015 (2015).

15. Donohoe, D. R. et al. The microbiome and butyrate regulate energy metabolism and autophagy in the mammalian colon. Cell metabolism 13, 517–526, doi:10.1016/j.cmet.2011.02.018 (2011).

16. Roediger, W. E. Role of anaerobic bacteria in the metabolic welfare of the colonic mucosa in man. Gut 21, 793–798, doi:10.1136/gut.21.9.793 (1980).

17. Cogliati, S., Enriquez, J. A. & Scorrano, L. Mitochondrial Cristae: Where Beauty Meets Functionality. Trends Biochem Sci 41, 261–273, doi:10.1016/j.tibs.2016.01.001 (2016).

18. Weber, J. S., Dummer, R., de Pril, V., Lebbe, C. & Hodi, F. S. Patterns of onset and resolution of immune-related adverse events of special interest with ipilimumab: detailed safety analysis from a phase 3 trial in patients with advanced melanoma. Cancer 119, 1675–1682, doi:10.1002/cncr.27969 (2013).

19. Wang, F., Yin, Q., Chen, L. & Davis, M. M. Bifidobacterium can mitigate intestinal immunopathology in the context of CTLA-4 blockade. Proceedings of the National Academy of Sciences of the United States of America 115, 157–161, doi:10.1073/pnas.1712901115 (2018).

20. Perez-Ruiz, E. et al. Prophylactic TNF blockade uncouples efficacy and toxicity in dual CTLA-4 and PD-1 immunotherapy. Nature 569, 428–432, doi:10.1038/s41586-019-1162-y (2019).

21. Mills, E. L. et al. Succinate Dehydrogenase Supports Metabolic Repurposing of Mitochondria to Drive Inflammatory Macrophages. Cell 167, 457–470.e413, doi:10.1016/j.cell.2016.08.064 (2016).

22. Bailis, W. et al. Distinct modes of mitochondrial metabolism uncouple T cell differentiation and function. Nature, doi:10.1038/s41586-019-1311-3 (2019).

23. Ghezzi, D. et al. SDHAF1, encoding a LYR complex-II specific assembly factor, is mutated in SDH-defective infantile leukoencephalopathy. Nat Genet 41, 654–656, doi:10.1038/ng.378 (2009).

24. Na, U. et al. The LYR factors SDHAF1 and SDHAF3 mediate maturation of the iron-sulfur subunit of succinate dehydrogenase. Cell metabolism 20, 253–266, doi:10.1016/j.cmet.2014.05.014 (2014).

25. Dotiwala, F. et al. Killer lymphocytes use granulysin, perforin and granzymes to kill intracellular parasites. Nature medicine 22, 210–216, doi:10.1038/nm.4023 (2016).

26. Harris, J. L., Peterson, E. P., Hudig, D., Thornberry, N. A. & Craik, C. S. Definition and redesign of the extended substrate specificity of granzyme B. The Journal of biological chemistry 273, 27364–27373, doi:10.1074/jbc.273.42.27364 (1998).

27. Wu, S. R. & Reddy, P. Tissue tolerance: a distinct concept to control acute GVHD severity. Blood 129, 1747–1752, doi:10.1182/blood-2016-09-740431 (2017).

28. Mehta, M. M., Weinberg, S. E. & Chandel, N. S. Mitochondrial control of immunity: beyond ATP. Nature reviews. Immunology 17, 608–620, doi:10.1038/nri.2017.66 (2017).

29. Vander Heiden, M. G., Cantley, L. C. & Thompson, C. B. Understanding the Warburg effect: the metabolic requirements of cell proliferation. Science (New York, N.Y.) 324, 1029–1033, doi:10.1126/science.1160809 (2009).

30. Chapman, N. M., Boothby, M. R. & Chi, H. Metabolic coordination of T cell quiescence and activation. Nature reviews. Immunology, doi:10.1038/s41577-019-0203-y (2019).

31. Sugiura, A. & Rathmell, J. C. Metabolic Barriers to T Cell Function in Tumors. Journal of immunology (Baltimore, Md.: 1950) 200, 400–407, doi:10.4049/jimmunol.1701041 (2018).

32. Chouchani, E. T. et al. Ischaemic accumulation of succinate controls reperfusion injury through mitochondrial ROS. Nature 515, 431–435, doi:10.1038/nature13909 (2014).

33. Caruso, R., Warner, N., Inohara, N. & Nunez, G. NOD1 and NOD2: signaling, host defense, and inflammatory disease. Immunity 41, 898–908, doi:10.1016/j.immuni.2014.12.010 (2014).

34. Graubert, T. A., DiPersio, J. F., Russell, J. H. & Ley, T. J. Perforin/granzyme-dependent and independent mechanisms are both important for the development of graft-versus-host disease after murine bone marrow transplantation. The Journal of clinical investigation 100, 904–911, doi:10.1172/jci119606 (1997).

35. Blazar, B. R., Taylor, P. A. & Vallera, D. A. CD4+ and CD8+ T cells each can utilize a perforin- dependent pathway to mediate lethal graft-versus-host disease in major histocompatibility complex-disparate recipients. Transplantation 64, 571–576, doi:10.1097/00007890-199708270-00004 (1997).

36. Jiang, Z., Podack, E. & Levy, R. B. Major histocompatibility complex-mismatched allogeneic bone marrow transplantation using perforin and/or Fas ligand double-defective CD4(+) donor T cells: involvement of cytotoxic function by donor lymphocytes prior to graft-versus-host disease pathogenesis. Blood 98, 390–397, doi:10.1182/blood.v98.2.390 (2001).

37. Luan, H. H. et al. GDF15 Is an Inflammation-Induced Central Mediator of Tissue Tolerance. Cell, doi:10.1016/j.cell.2019.07.033 (2019).

38. Gopalakrishnan, V. et al. Gut microbiome modulates response to anti-PD-1 immunotherapy in melanoma patients. *Science (New York*, N.Y.) 359, 97–103, doi:10.1126/science.aan4236 (2018).

39. Matson, V. et al. The commensal microbiome is associated with anti-PD-1 efficacy in metastatic melanoma patients. *Science (New York*, N.Y.) 359, 104–108, doi:10.1126/science.aao3290 (2018).

40. Routy, B. et al. Gut microbiome influences efficacy of PD-1-based immunotherapy against epithelial tumors. *Science (New York*, N.Y.) 359, 91–97, doi:10.1126/science.aan3706 (2018).

41. Chung, K. Y. et al. Phase II study of the anti-cytotoxic T-lymphocyte-associated antigen 4 monoclonal antibody, tremelimumab, in patients with refractory metastatic colorectal cancer. Journal of clinical oncology: official journal of the American Society of Clinical Oncology 28, 3485–3490, doi:10.1200/jco.2010.28.3994 (2010).

42. Le, D. T. et al. PD-1 Blockade in Tumors with Mismatch-Repair Deficiency. The New England journal of medicine 372, 2509–2520, doi:10.1056/NEJMoa1500596 (2015).

43. Yardeni, T. et al. Host mitochondria influence gut microbiome diversity: A role for ROS. Science signaling 12, doi:10.1126/scisignal.aaw3159 (2019).

44. Eriguchi, Y. et al. Graft-versus-host disease disrupts intestinal microbial ecology by inhibiting Paneth cell production of alpha-defensins. Blood 120, 223–231, doi:10.1182/blood-2011-12-401166 (2012).

45. Jenq, R. R. et al. Intestinal Blautia Is Associated with Reduced Death from Graft-versus-Host Disease. Biol Blood Marrow Transplant 21, 1373–1383, doi:10.1016/j.bbmt.2015.04.016 (2015).

46. Jenq, R. R. et al. Regulation of intestinal inflammation by microbiota following allogeneic bone marrow transplantation. The Journal of experimental medicine 209, 903–911, doi:10.1084/jem.20112408 (2012).

47. Shono, Y. et al. Increased GVHD-related mortality with broad-spectrum antibiotic use after allogeneic hematopoietic stem cell transplantation in human patients and mice. Sci Transl Med 8, 339ra371, doi:10.1126/scitranslmed.aaf2311 (2016).

48. Peled, J. U. et al. Intestinal Microbiota and Relapse After Hematopoietic-Cell Transplantation. Journal of clinical oncology: official journal of the American Society of Clinical Oncology 35, 1650–1659, doi:10.1200/jco.2016.70.3348 (2017).

49. Atarashi, K. et al. Induction of colonic regulatory T cells by indigenous Clostridium species. *Science (New York*, N.Y.) 331, 337–341, doi:10.1126/science.1198469 (2011).

50. Atarashi, K. et al. Treg induction by a rationally selected mixture of Clostridia strains from the human microbiota. Nature 500, 232–236, doi:10.1038/nature12331 (2013).

51. Furusawa, Y. et al. Commensal microbe-derived butyrate induces the differentiation of colonic regulatory T cells. Nature 504, 446–450, doi:10.1038/nature12721 (2013).

52. Fujiwara, H. et al. Microbial metabolite sensor GPR43 controls severity of experimental GVHD. Nature communications 9, 3674, doi:10.1038/s41467-018-06048-w (2018).

53. Kakihana, K. et al. Fecal microbiota transplantation for patients with steroid-resistant acute graft-versus-host disease of the gut. Blood 128, 2083–2088, doi:10.1182/blood-2016-05-717652 (2016).

54. DeFilipp, Z. et al. Third-party fecal microbiota transplantation following allo-HCT reconstitutes microbiome diversity. Blood advances 2, 745–753, doi:10.1182/bloodadvances.2018017731 (2018).

55. Biton, M. et al. T Helper Cell Cytokines Modulate Intestinal Stem Cell Renewal and Differentiation. Cell 175, 1307–1320.e1322, doi:10.1016/j.cell.2018.10.008 (2018).

56. Cooke, K. R. et al. Tumor necrosis factor- alpha production to lipopolysaccharide stimulation by donor cells predicts the severity of experimental acute graft-versus-host disease. The Journal of clinical investigation 102, 1882–1891, doi:10.1172/jci4285 (1998).

57. Hill, G. R. et al. Interleukin-11 promotes T cell polarization and prevents acute graft-versus-host disease after allogeneic bone marrow transplantation. The Journal of clinical investigation 102, 115–123, doi:10.1172/jci3132 (1998).

58. Sas, K. M. et al. Tissue-specific metabolic reprogramming drives nutrient flux in diabetic complications. JCI insight 1, e86976, doi:10.1172/jci.insight.86976 (2016).

59. Mathew, A. V. et al. Impaired Amino Acid and TCA Metabolism and Cardiovascular Autonomic Neuropathy Progression in Type 1 Diabetes. Diabetes 68, 2035–2044, doi:10.2337/db19-0145 (2019).

60. Mathew, A. V., Seymour, E. M., Byun, J., Pennathur, S. & Hummel, S. L. Altered Metabolic Profile With Sodium-Restricted Dietary Approaches to Stop Hypertension Diet in Hypertensive Heart Failure With Preserved Ejection Fraction. Journal of cardiac failure 21, 963–967, doi:10.1016/j.cardfail.2015.10.003 (2015).

61. Halbrook, C. J. et al. Macrophage-Released Pyrimidines Inhibit Gemcitabine Therapy in Pancreatic Cancer. Cell metabolism 29, 1390–1399.e1396, doi:10.1016/j.cmet.2019.02.001 (2019).

62. Martin, S. J. et al. The cytotoxic cell protease granzyme B initiates apoptosis in a cell-free system by proteolytic processing and activation of the ICE/CED-3 family protease, CPP32, via a novel two-step mechanism. The EMBO journal 15, 2407-2416 (1996).

63. West, A. P. et al. Mitochondrial DNA stress primes the antiviral innate immune response. Nature 520, 553–557, doi:10.1038/nature14156 (2015).

