## Supplementary material for "Mitochondrial Complex II In Intestinal Epithelial Cells Regulates T-cell Mediated Immunopathology": Supplemetary Figure

### Supplementary Figure 1

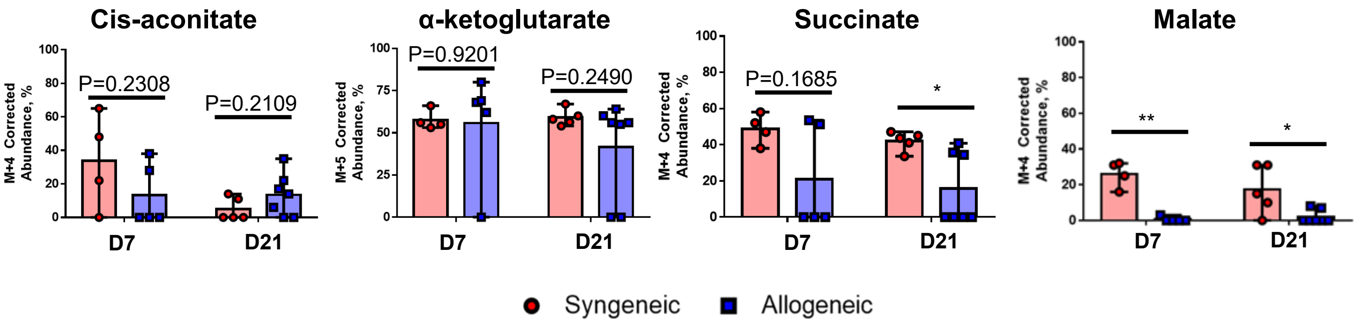

### Supplementary Figure 2

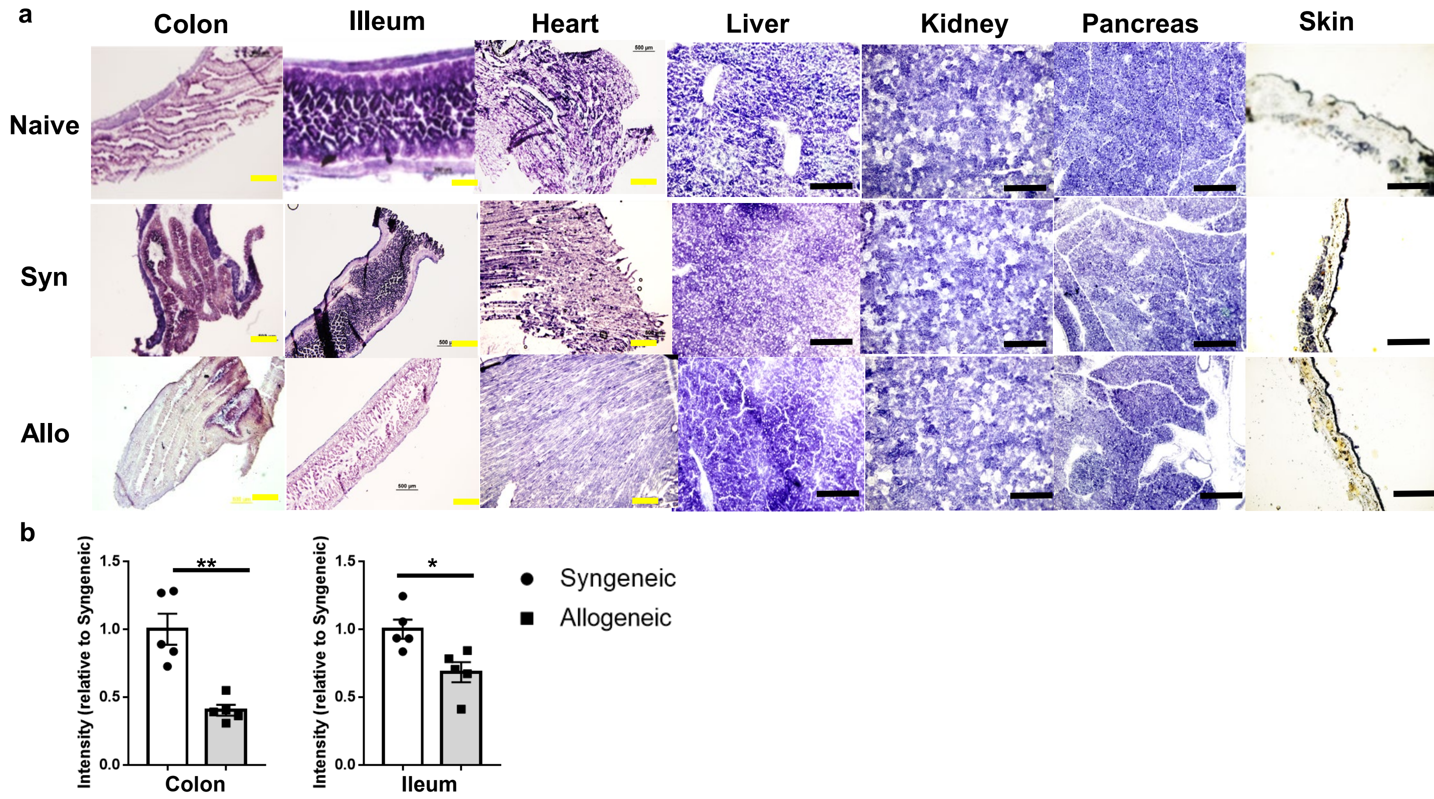

### Supplementary Figure 3

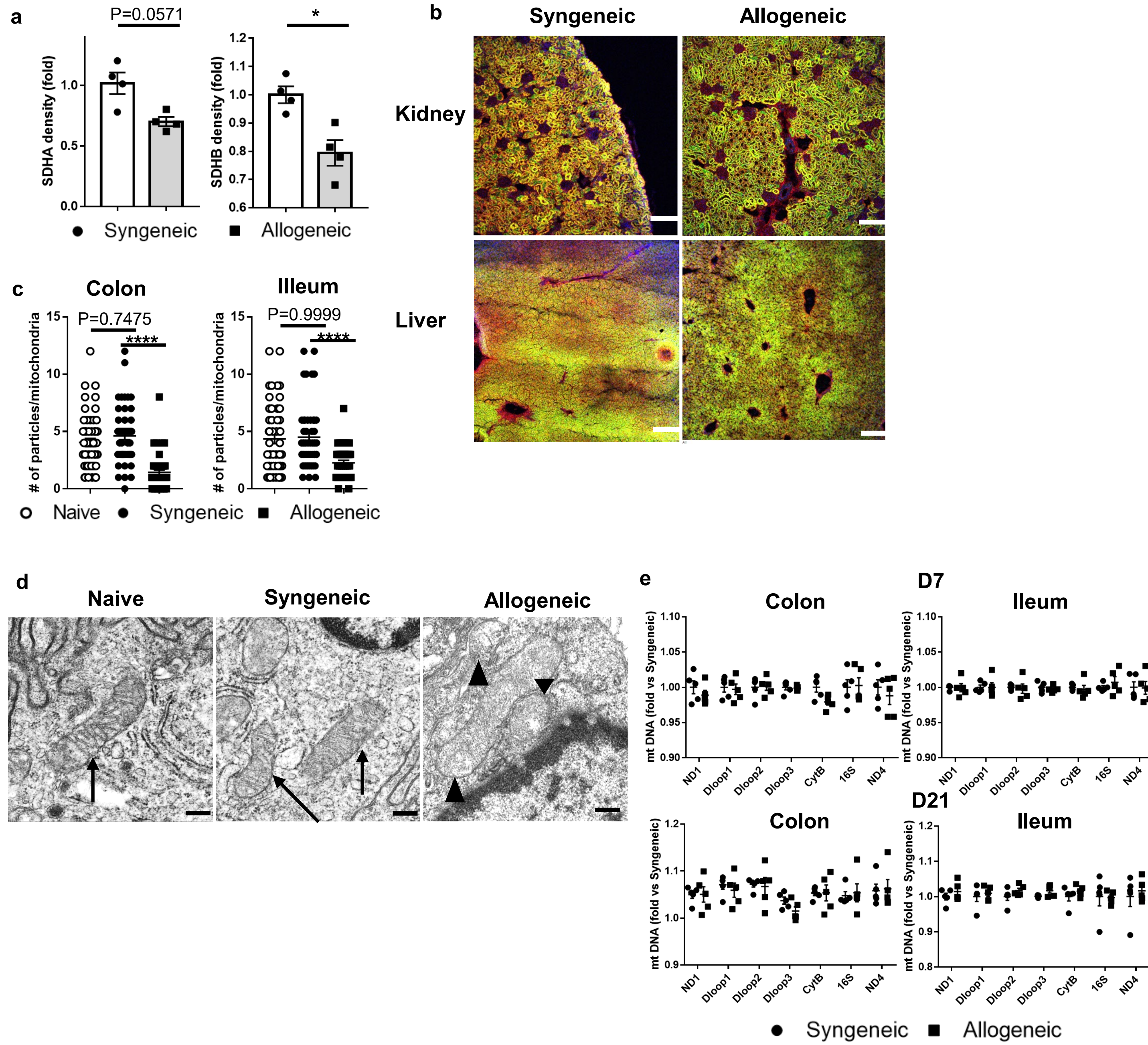

### Supplementary Figure 4

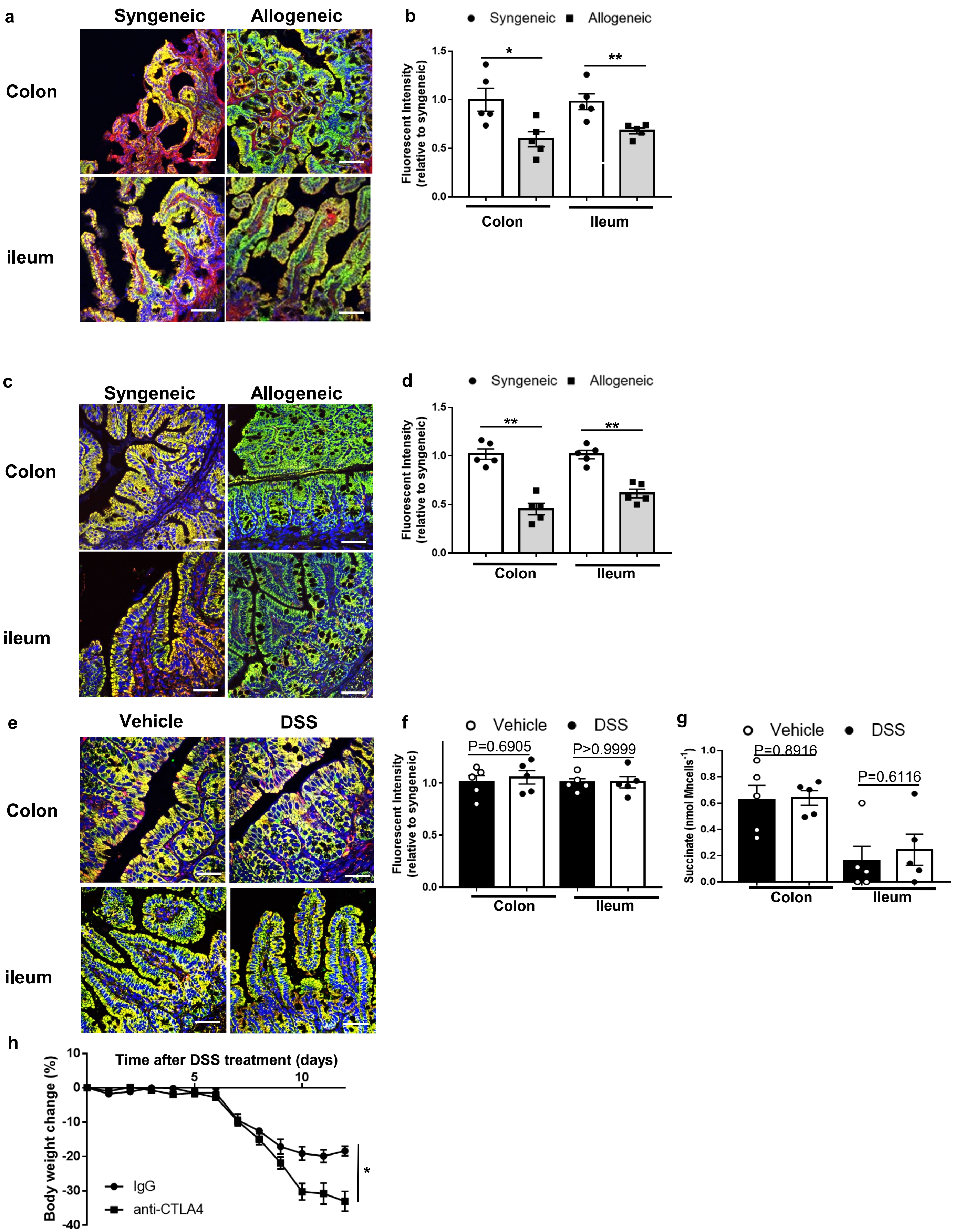

### Supplementary Figure 5

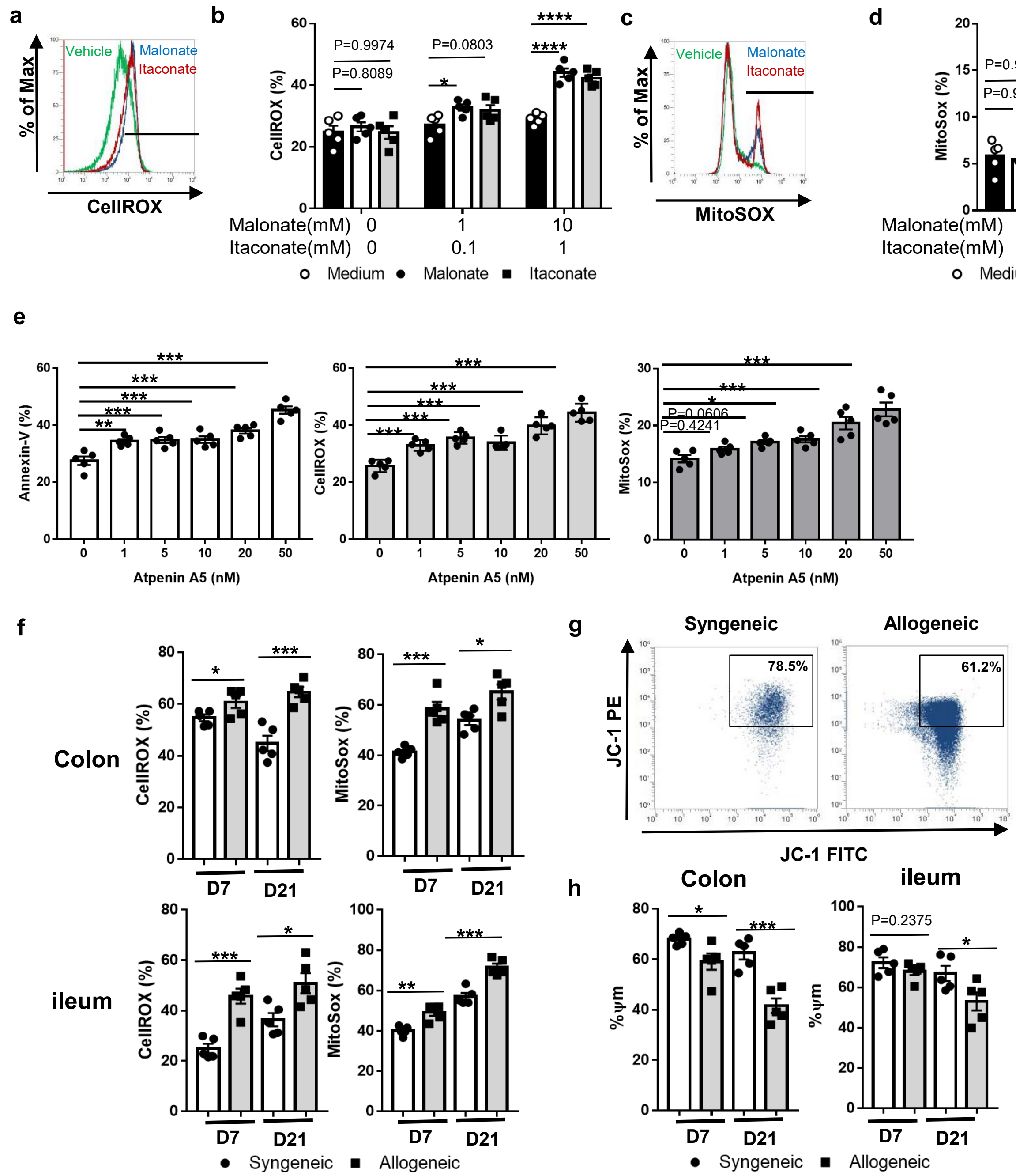

### Supplementary Figure 6

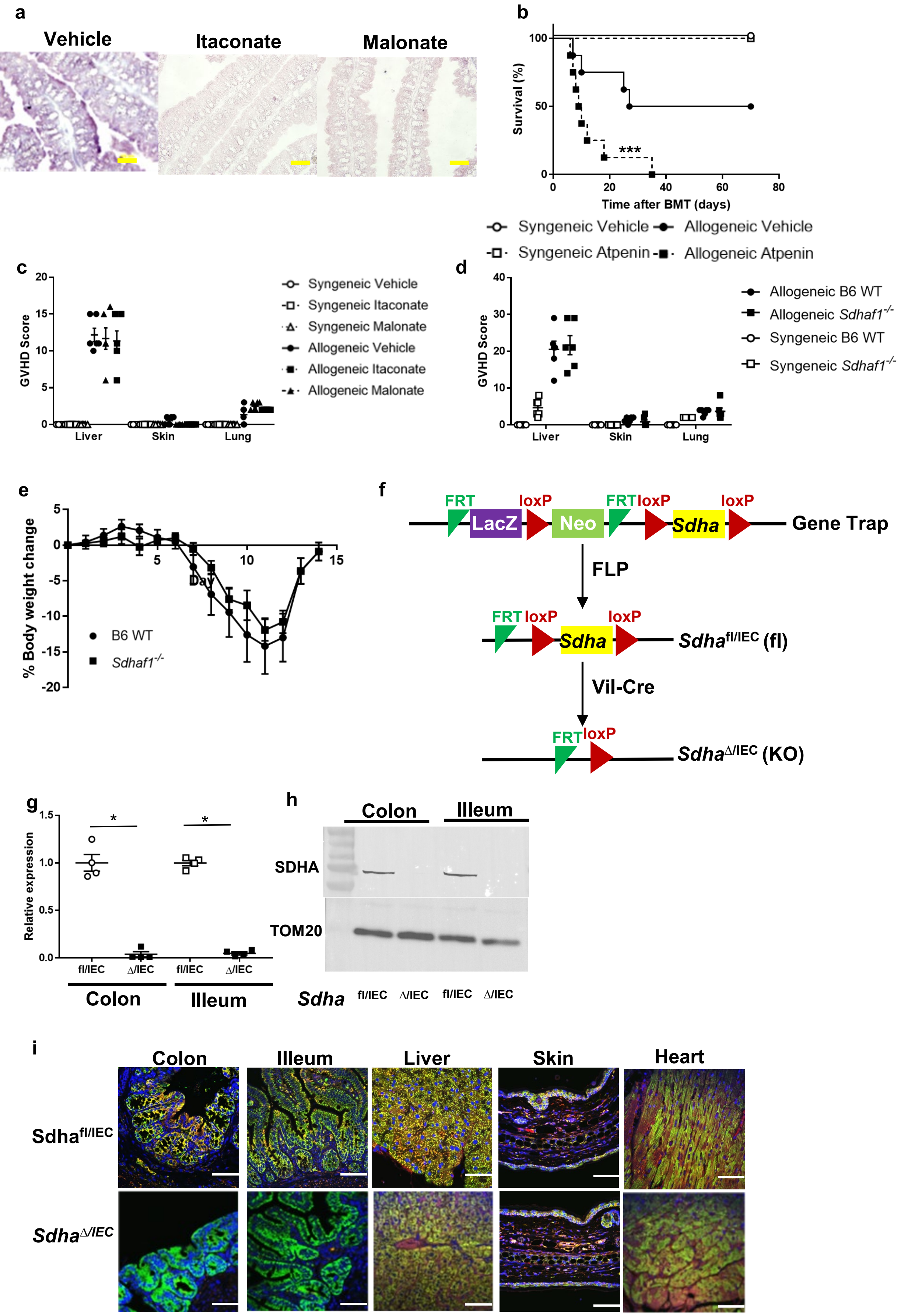

### Supplementary Figure 7

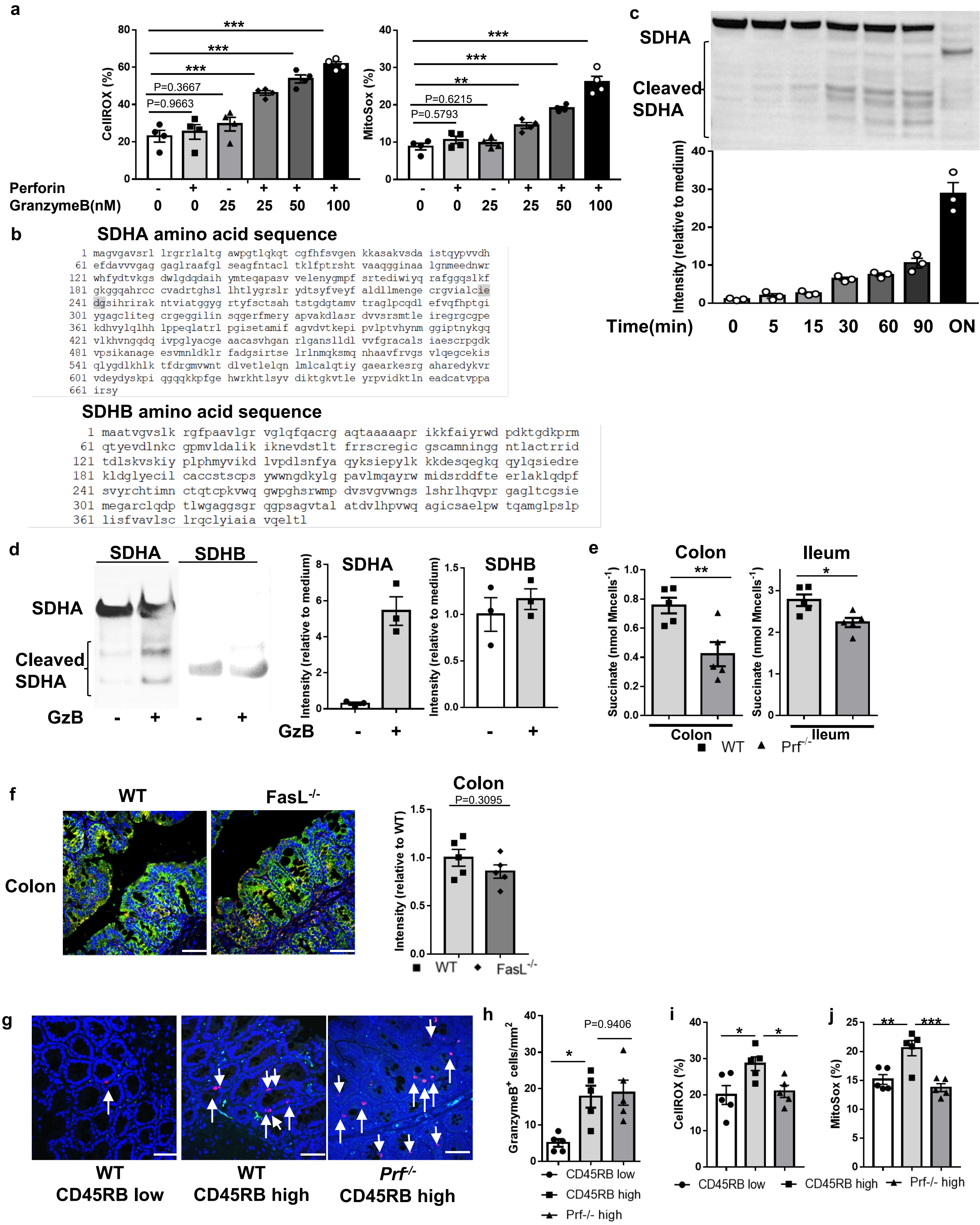

### Supplementary Figure 8

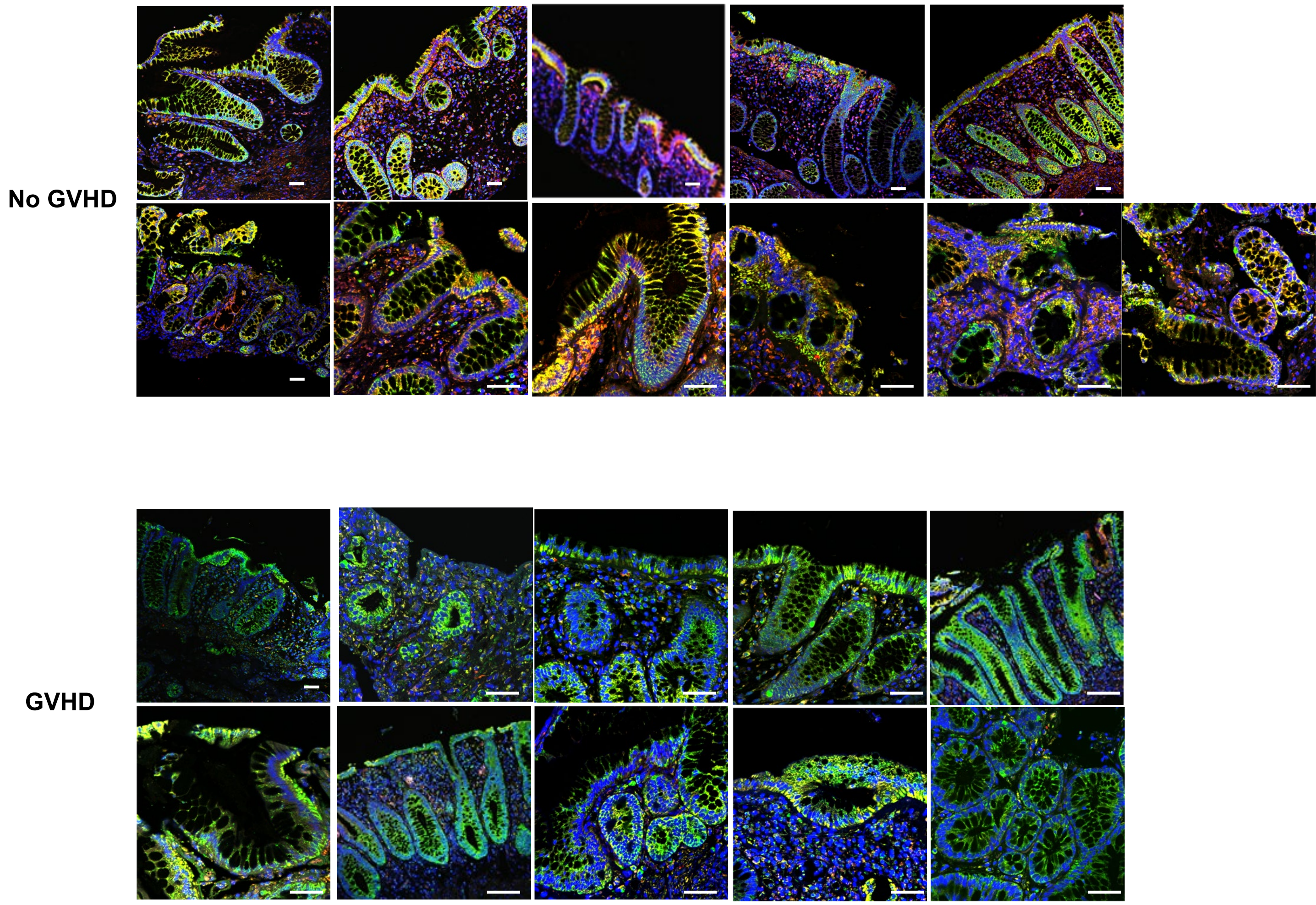
